# Combinatorial community coalescence in early tomato assembly reveals a rhizosphere attractor in composition and abundance architecture

**DOI:** 10.64898/2026.07.05.736546

**Authors:** Rubén Chaboy-Cansado, Paula Cobeta, Gabriel Roscales, Alberto Rastrojo, Daniel Aguirre de Cárcer

## Abstract

The rhizosphere microbiome plays fundamental roles in plant health and productivity, yet the ecological rules governing microbiome assembly remain poorly understood. Here, we investigated early rhizosphere community assembly in tomato using a replicated combinatorial community coalescence framework, in which seven distinct natural bacterial communities were inoculated individually and in all possible pairwise and triplet combinations.

Single-inoculum communities clustered according to inoculum identity, indicating a strong effect of source community composition on assembly trajectories. However, when all communities were analyzed jointly, samples formed a continuous compositional landscape with no clear evidence of discrete community states. Despite major differences in source community composition, rhizosphere communities consistently converged toward a highly similar uneven rank–abundance structure, with two ASVs accounting for 50% and a median of nineteen ASVs for 90% of total abundance. While assembly was dominated by a very small number of *Pseudomonas* ASVs, limited evidence of alternative dominant states was observed. Increasing inoculum complexity did not increase stochasticity but instead promoted stronger convergence toward a global rhizosphere compositional centroid. Moreover, dominance hierarchies emerging from community coalescence closely mirrored the distance of source communities to this centroid. The two most dominant communities originated from orchard soils, suggesting that historical contingency and prior adaptation to horticultural crop rhizospheres may influence competitive success.

Together, these results are consistent with the existence of a canonical rhizosphere attractor in composition and abundance architecture, with patterns consistent with assembly occurring under a limited number of dominant ecological niches imposed by the tomato rhizosphere.

## INTRODUCTION

The rhizosphere, defined as the narrow soil zone directly affected by plant roots, hosts one of the most diverse and metabolically active microbial ecosystems on Earth ^1^. Within this highly dynamic environment, plants continuously modulate the surrounding microbial community through the secretion of root exudates, mucilage, and cellular debris, thereby favoring the establishment of specific microbial assemblages ^2,3^. In return, rhizosphere microorganisms can enhance plant performance through multiple mechanisms, including increasing nutrient availability and phytohormone production ^4^, improving tolerance to abiotic stresses ^5,6^, and suppressing pathogens via competition or antagonistic interactions ^7^.

Understanding the ecological and mechanistic drivers of rhizosphere community dynamics has become increasingly relevant. This is particularly important in the ongoing shift toward more sustainable agricultural systems, where microbiome-based strategies and products are expected to play a major role. Progress in this area depends on a deeper understanding of rhizosphere microbiome assembly and how microbial community composition relates to ecosystem function. Despite major advances in rhizosphere microbiome research, predicting how microbial communities assemble around plant roots remains a central challenge. Rhizosphere communities emerge from highly diverse soil microbial pools through a combination of dispersal, host-mediated selection, microbial interactions, stochastic colonization, and temporal succession during plant development ^8–10^. Although these processes can generate substantial taxonomic variation, growing evidence suggests that rhizosphere assembly may nevertheless follow partially predictable ecological rules. Across diverse plant systems, root-associated microbiomes often display recurrent compositional patterns and consistent enrichment of particular bacterial lineages, indicating that plant-associated selection can constrain assembly toward a limited set of successful colonizers ^6,11^.

A major unresolved question is whether rhizosphere microbiome assembly tends to converge toward common community configurations or instead allows multiple alternative assembly outcomes. In many microbial ecosystems, community structure is characterized by highly uneven abundance distributions, typically composed of a small number of dominant taxa and a long tail of low-abundance organisms. Such recurrent community structures suggest that strong ecological filtering and competitive interactions may drive communities toward constrained and potentially predictable states. However, historical contingency, stochasticity, and priority effects may also generate alternative trajectories, even under similar environmental conditions. Distinguishing between convergence toward common attractors and divergence into alternative stable states remains a major challenge for understanding microbiome assembly dynamics ^12,13^. Here, we use the term rhizosphere attractor operationally to refer to a recurrent region of community-composition space and abundance organization toward which independently assembled communities converge under the same host and environmental conditions.

Importantly, most rhizosphere microbiome studies have examined assembly from a single environmental microbial pool. However, in natural and agricultural systems, roots are frequently exposed to multiple microbial communities simultaneously or sequentially through soil mixing, organic amendments, microbial inoculants, agricultural practices, and root–root interactions. This raises a fundamental ecological question: when multiple microbial communities encounter the same host environment, do they assemble into reproducible rhizosphere states or generate divergent community trajectories?

An emerging framework to address this question is community coalescence, defined as the encounter and mixing of previously isolated microbial ^14^. Community coalescence is increasingly recognized as a pervasive process in microbial ecology and a major driver of microbiome assembly in natural and managed systems. Importantly, coalescence outcomes are often strongly non-additive, ranging from near-complete mixing to pronounced asymmetrical dominance by one of the original communities. Theoretical and experimental studies suggest that these outcomes can be shaped by propagule pressure, priority effects, competitive asymmetries, and emergent community-level cohesion, whereby taxa from the same source community facilitate each other’s persistence during coalescence ^13,15^. Community coalescence therefore provides a powerful conceptual and experimental framework to investigate microbiome predictability, convergence, and contingency.

Tomato (*Solanum lycopersicum*) is both one of the most economically important crop species and a major model system for plant–microbiome research ^16^. Previous studies have provided valuable insights into tomato-associated microbial communities across plant compartments and into the role of host genetics in microbiome assembly ^17–19^. However, the ecological dynamics governing early rhizosphere assembly in tomato, particularly under conditions involving multiple competing microbial source communities, remain poorly understood.

Here, we investigated early rhizosphere microbiome assembly in tomato using a controlled experimental framework designed to examine both microbiome predictability and community coalescence outcomes. We inoculated tomato plants with seven distinct natural bacterial communities originating from different soils, both individually and in all possible pairwise and triplet combinations, thereby generating a broad range of controlled coalescence events. Pairwise combinations were used to characterize directional coalescence outcomes, whereas the complete dataset including single, pairwise and triplet inocula was used to evaluate global assembly landscapes and community phenotype effects. To minimize host genetic effects, a single tomato genotype was used throughout, and plants were grown under sterile conditions in vermiculite-based microcosms. After three weeks of growth, we characterized rhizosphere community composition and quantified bacterial load and plant performance.

## MATERIALS AND METHODS

### Experimental design

Seven distinct natural bacterial communities were inoculated separately onto tomato plants and in all possible combinations of two and three communities, with three replicates per treatment. After three weeks of plant growth rhizospheres were destructively sampled.

### Planting and inoculation

The microbial fraction (MF) was isolated from different natural soils (Suppl. Table 1) and titrated as previously described ^20^. The communities were labeled A, C, D, E, F, G, and H following the nomenclature established in previous experiments. Previous studies have demonstrated that plant genotype can influence rhizosphere composition ^21,22^. To minimize this potential confounding factor, seeds from a single genetically homogeneous tomato variety, Salinas F1 (CapGen Seeds), were used throughout the experiment. Seeds were surface-sterilized before germination in the dark in water-agar plates ^20^. Seedlings with roots approximately 2–3 mm long were selected and transferred into sterile 40 mL polypropylene tubes containing 3 g of pre-sterilized vermiculite. Immediately after transplantation, each seedling was watered with 15 mL of Gamborg B5 plant culture medium (Duchefa Biochemie) previously inoculated with the corresponding bacterial community. Each plant received an inoculum containing 9,2 × 10^4^ bacteria from the MFs or their mixtures combined in equal proportions.

Plants were grown in sterile greenhouses under a 16 h light (24 °C) / 8 h dark (16 °C) photoperiod. Greenhouses were sterilized by washing all surfaces with 70% ethanol, sealing ventilation openings with autoclaved cotton, and exposing internal surfaces to ultraviolet light for 5 min.

### Plant processing

The greenhouses were opened under a laminar-flow hood to recover the tubes containing the plants. Shoots were separated from roots using sterile forceps, and as much vermiculite as possible attached to the roots was carefully removed using the same forceps. Shoots were subsequently dried at 65 °C for 48 h in an oven (JP Selecta, model 200) to remove moisture and determine dry weight using a precision balance. Roots were transferred into sterile 40 mL polypropylene tubes containing 2 mL of PBS to obtain rhizosphere microbial fractions (hereafter RMFs), and samples were kept on ice. Once all roots had been transferred to their corresponding tubes, samples were vortexed for 10 min at 1490 rpm and 4 °C using a Multi Reax mixer (Heidolph). After vortexing, 1 mL of each RMF was transferred into a sterile 1.5 mL polypropylene tube. Aliquots from each sample were then distributed into sterile 0.2 mL polypropylene tube strips for subsequent DNA extraction and titration, which was carried out as previously described ^20^.

### Community profiling

Community DNA was extracted from each RMF following the microvolume alkaline lysis protocol described by Bramucci et al. (2021) ^23^. The bacterial 16S rRNA gene was amplified using primers 799F and 1193R, targeting the V5–V7 hypervariable region and incorporating Illumina sequencing adapters separated by 0–7 random nucleotides. This design introduces a phase shift during sequencing, preventing signal overlap from highly similar amplicons and substantially improving sequencing performance ^24^. PCR reactions were performed in a final volume of 25 µL containing 4 µL of template DNA, 0.1 µM of each primer, 0.4 mM dNTPs, and 0.5 U of Q5 High-Fidelity DNA Polymerase (NEB). Thermocycling conditions consisted of an initial denaturation at 95 °C for 30 s, followed by 25 cycles of 95 °C for 10 s, 55 °C for 30 s, and 72 °C for 30 s, with a final extension at 72 °C for 2 min. PCR products were then diluted 1:10 in 20 µL of nuclease-free water to reduce carryover of spurious amplicons and residual oligonucleotides from the first amplification. A second PCR of 10 cycles was subsequently performed under identical cycling conditions using 3 µL of diluted PCR product as template and 4 µL of 5 µM secondary primers containing Illumina i5 and i7 adapter sequences, 10-nt barcodes, and Illumina sequencing primer sites. Within each PCR batch, samples shared a common barcode at one end and carried unique barcodes at the other, enabling unambiguous assignment of sequencing reads to individual samples.

Amplicon libraries were evaluated on 1.5% agarose gels. Band intensities were visually assessed within each batch to pool approximately equimolar quantities of each sample. Pooled libraries were concentrated using Pellet Paint® NF Co-Precipitant (Merck) according to the manufacturer’s instructions, followed by size selection using horizontal agarose gel electrophoresis and purification with the Speedtools PCR Clean-Up Kit (Biotools). Final pooled amplicon libraries were sequenced on an Illumina NextSeq platform using a NextSeq1000/2000 P1 flow cell with 2 × 300 bp paired-end sequencing.

### Sequence processing and analyses

Initial sequence processing was carried out using the *DADA2* package in R ^25^, following the standard workflow for error modelling, paired-end read merging, chimera removal, taxonomic assignment against the SILVA training database (v123) ^26^, and removal of remaining eukaryotic sequences, including reads derived from mitochondria and chloroplasts. To standardize sequencing depth across samples, rarefaction was performed at a common threshold defined by the natural break in sequence-count distribution ^27^ (Suppl. Fig. 1), and samples with fewer than 23,394 sequences were excluded. For phylogeny-based analyses, amplicon sequence variants (ASVs) were mapped to the Greengenes2 99% representative sequence database and associated phylogenetic tree ^28^ using BLAST. Faith’s phylogenetic diversity was estimated using the *pd* function from the *picante* package (v1.8.2) ^29^.

### Coalescence analysis

Data subsets were generated containing samples from each pairwise community combination and the corresponding individual communities. Community composition within each subset was visualized by non-metric multidimensional scaling (NMDS) ordinations based on Bray-Curtis and UniFrac distance matrices using the *ordinate* function from the *phyloseq* package (v1.52.0). Pairwise dissimilarity matrices were additionally calculated using the *distance* function from the same package. From these matrices, dissimilarities were extracted for: i) comparisons among samples within each individual community (intra-group comparisons), and ii) comparisons between samples from each individual community and the corresponding mixed community (inter-group comparisons). Thus, for each subset and distance metric, four comparisons were obtained (two intra-group and two inter-group) and visualized using boxplots. To visualize ASV overlap between individual communities and their corresponding mixtures, Euler diagrams were generated using the *euler* function from the *eulerr* package (v7.0.4). Relative ASV abundances were further represented for each subset using rainbow plots, distinguishing ASVs shared among individual communities and mixtures from those exclusive to a specific community. These plots were generated using custom functions developed by our research group (github.com/silvtal/rainbow_function).

### Community load and plant dry weight statistical analyses

The effects of ASV richness and phylogenetic diversity (Faith’s index) on bacterial load and plant dry weight were evaluated. Because Faith’s phylogenetic diversity is calculated as the sum of branch lengths across taxa, it inherently includes a richness component, potentially leading to collinearity and complicating the interpretation of their individual effects. To assess the relationship between Richness and Diversity, the Pearson correlation coefficient between ASV richness and Faith’s diversity was assessed using the *cor* function in R.

To obtain the fraction of diversity independent of ASV richness for subsequent analyses, residuals from a simple linear model of diversity against richness were extracted. The effects of richness and this independent diversity component on plant dry weight were evaluated using a linear model. For bacterial load, values were log10-transformed prior to analysis to improve normality. In both cases, effect significance was assessed using Type II ANOVA with the *Anova* function from the *car* package (v3.1-3). Model heteroscedasticity was evaluated using *bptest* from the *lmtest* package (v0.9-40).

To identify ASVs associated with dry weight, putative source sets were first defined for each ASV based on the single-inoculum communities. An ASV was considered associated with a given single inoculum when its mean relative abundance across replicate samples of that inoculum was at least 0.1%. Eligible samples for each ASV were then defined as all samples whose inoculum combination contained at least one of the ASV source components. ASVs were retained for analysis when they reached a relative abundance of at least 0.1% in at least 10% of their eligible samples. For each retained ASV, associations between relative abundance and dry weight were evaluated using Spearman rank correlations calculated with the *cor.test* function from the *stats* package in R. P-values were adjusted across ASVs using the Benjamini–Hochberg false discovery rate (FDR) procedure implemented with the *p.adjust* function from the *stats* package.

To assess ASV associations with dry weight independently of the average effect of inoculum composition, dry weight was additionally modelled as a function of inoculum combination using the *lm* function from the *stats* package in R. Residuals from this model were extracted and used as corrected dry weight values representing deviations from the mean expected dry weight for each inoculum combination. Spearman rank correlations were then recalculated between ASV relative abundance and residual dry weight using the same eligibility criteria and statistical procedures described above.

To evaluate biomass-associated patterns at broader taxonomic resolution, ASV abundances were collapsed at genus and family levels. Taxa with relative abundance ≥0.1% in at least 10 samples were retained for analysis. Associations between taxon relative abundance and dry weight were evaluated using Spearman rank correlations as described above. Analyses were performed using both raw dry weight values and residual dry weight values obtained after correcting for inoculum- combination effects.

To evaluate whether plant performance was associated with global community structure, Bray– Curtis distances among samples were calculated using the *vegdist* function from the *vegan* package (v2.6-10). A global compositional centroid was defined as the mean relative abundance profile across all samples, and each sample’s distance to this centroid was calculated from the resulting Bray–Curtis distance matrix and used as a proxy for proximity to the global community attractor. Associations between centroid distance and dry weight were assessed using Spearman rank correlations as described above.

To further evaluate selected candidate biomass-associated ASVs, a targeted exploratory analysis was performed for ASV_106 and ASV_117. Samples were classified according to the presence or absence of each ASV using a relative abundance threshold of 0.1%, resulting in four groups: neither ASV present, ASV_106 only, ASV_117 only, and both ASVs present. Dry weight distributions were compared across groups using descriptive statistics and Wilcoxon rank-sum tests implemented with the *wilcox.test* function from the *stats* package.

### Community assembly landscape and constraint analyses

For analyses focused on dominant and reproducibly detected taxa, a filtered ASV table was generated by retaining only ASVs reaching at least 0.1% relative abundance in at least two samples. This filtering step removed extremely rare taxa while preserving ASVs with evidence of reproducible ecological establishment.

Global patterns of community composition were explored using non-metric multidimensional scaling (NMDS) based on Bray–Curtis dissimilarities. Ordinations were performed using the *ordinate* function from the *phyloseq* package (v1.52.0), using 100 random starts to improve convergence. Two ordinations were generated: one including all rhizosphere communities (single, pair, and triplet inocula), and a second restricted to single-inoculum communities only.

Rank–abundance distributions were used to characterise global community structure and dominance patterns. To summarise global patterns across samples, rank-wise abundance envelopes were calculated using the median, interquartile range (25th–75th percentile), and broader distribution range (5th–95th percentile) at each rank position. These envelopes were generated for all samples combined and separately for single, pair, and triplet inocula. An additional analysis restricted to ASVs exceeding 1% relative abundance was performed to examine dominance patterns among abundant taxa. To quantify dominance strength, cumulative rank–abundance curves were calculated for each sample, and the number of ASVs required to explain 50% and 90% of total community relative abundance was extracted and summarised across samples.

To evaluate assembly reproducibility, pairwise Pearson correlations were calculated between all pairs of samples using ASV relative abundance profiles using the *cor* function from the *stats* package in R. Sample pairs were classified as intra-combination when both samples belonged to the same inoculum combination, or inter-combination when samples belonged to different inoculum combinations.

To quantify convergence toward a common rhizosphere compositional state, a global compositional centroid was defined as the mean relative abundance profile across all samples. Bray–Curtis distances between each sample and the centroid were calculated using the *vegdist* function from the *vegan* package (v2.6-10), and these distances were used as a proxy for distance to the global rhizosphere attractor.

To characterize taxon-specific assembly behavior, ASV-level assembly metrics were calculated for filtered ASVs. For each ASV, its source set was defined as the set of single inocula in which the ASV was detected. An inoculum combination was considered eligible for a given ASV when it contained at least one member of the ASV source set. Three ASV-level assembly metrics were then calculated. Conditional persistence quantified how often an ASV established when ecologically possible and was calculated as the proportion of eligible mixtures in which the ASV was detected. Retention quantified how well an ASV maintained abundance after mixing relative to its best-performing source inoculum and was calculated as the mean log2 ratio between ASV relative abundance in each eligible mixture and its maximum abundance across relevant source single inocula, using a pseudo count of 10⁻⁶ to avoid division by zero. Context dependence quantified sensitivity to assembly context and was calculated as the standard deviation of retention values across eligible mixtures. Source breadth was defined as the number of single inocula in which the ASV was detected and used as a proxy for ecological generalism.

All scripts and 16S rRNA gene dataset are available at https://github.com/microenvgen/Coalescence and the European Nucleotide Archive (XXXXX), respectively. During the preparation of this manuscript, the authors used generative AI tools (ChatGPT, OpenAI) for language editing, translation support, and assistance with code troubleshooting and scripting. All generated suggestions were critically reviewed, validated, and implemented by the authors. The authors take full responsibility for the final content, analyses, and conclusions.

## RESULTS

The rhizosphere microbiome of our experimental system was dominated by bacterial families typically associated with this type of ecosystem, such as Pseudomonadaceae, Xanthomonadaceae, and Comamonadaceae (Figure 1). To evaluate the outcome of subjecting distinct microbial communities to coalescence events, we focused on the rhizosphere communities arising from single and pairwise inoculations. To determine whether the communities resulting from coalescence represented a mixture of the original communities or, alternatively, whether one community dominated the other, several complementary analyses were performed. First, non-metric multidimensional scaling (NMDS) ordinations based on Bray-Curtis and UniFrac distances were used to assess compositional similarity among communities. In addition, Bray-Curtis and UniFrac distances were represented using boxplots for both intra-group comparisons (among samples belonging to the same individual community) and inter-group comparisons (between samples from individual communities and their corresponding coalesced community). Euler diagrams were further used to quantify the ASVs contributed by each individual community to the coalesced community, whereas comparative ASV abundance plots were used to identify which community contributed the most abundant ASVs to the final mixture. It should be noted that interpretation of these analyses remained partly subjective and, in some cases, it was difficult to conclusively determine whether a dominant community emerged.

**Figure 1.**
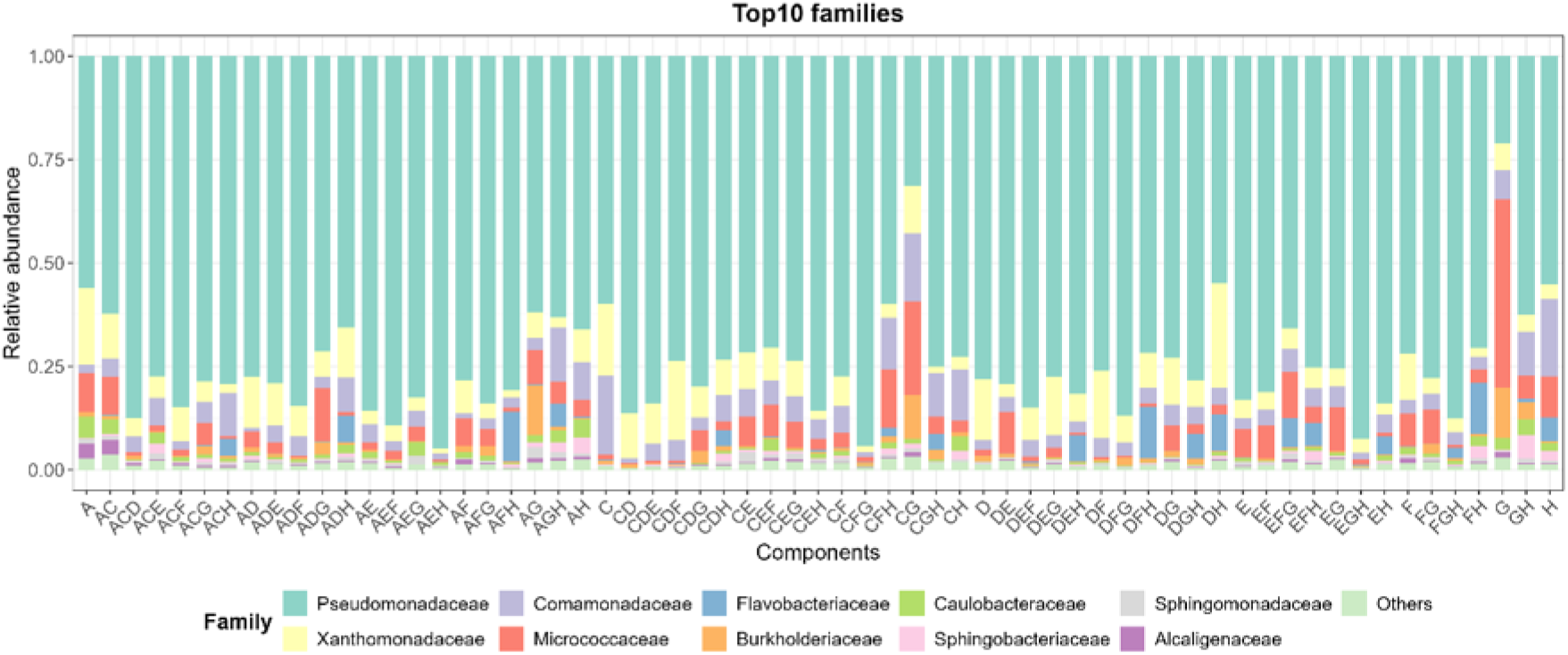
Relative abundance of the ten most abundant bacterial families across the dataset in the studied samples. The Y-axis represents the distribution of mean relative abundance values for each family across the different microbial communities studied (individual communities or community combinations; X-axis).

Figure 2 and 3 illustrate representative examples of these analyses (dominance and no dominance, respectively), whereas the remaining coalescence events are shown in the supplementary material (Fig. S2–S21). After analyzing all coalescence experiments, a directed network was constructed to summarize the outcomes of these comparisons (Fig. 4). As shown by the results, Community E dominated all other communities, followed by Community D, which exhibited the second highest dominance rate. In contrast, Communities C and G did not dominate any other community, and no clear dominance pattern was observed in the coalescence event between them. When diversity metrics were examined (Supplementary table 2), no clear relationship was observed between community diversity and dominance rate during coalescence events. However, when considering the soil origin of each community, an exploratory association with soil-use history emerged (Figure 4. Supplementary table 1-2). The communities with the highest dominance rates originated from orchard garden soils (Community E from Ruiseñada and Community D from Vigo), whereas communities with the lowest dominance rates originated from dryland agricultural soil (Community G from Badajoz) and forest soil (Community C from Valdelatas). It is worth noting that Community A also originated from an orchard garden soil, although it showed lower dominance than communities originating from ruderal soils (Communities F and H). However, given the small number of source communities and the unbalanced representation of soil-use categories, this pattern should be regarded as exploratory rather than as evidence for a general effect of soil-use history on coalescence outcomes.

**Figure 2.**
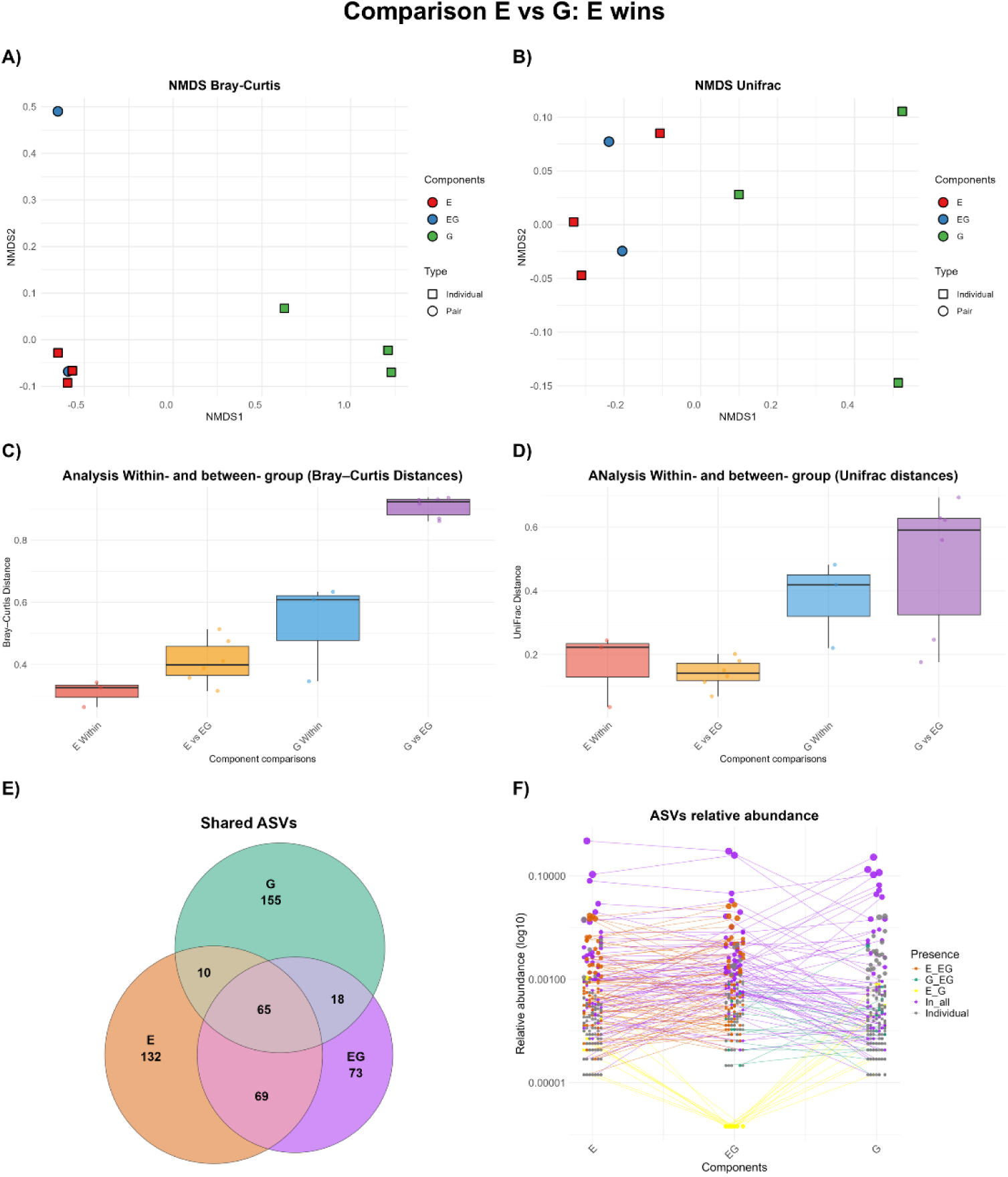
Representation of the different analyses performed to characterize the microbial communities resulting from community coalescence (EvsG). A) NMDS ordination based on Bray– Curtis distances of the microbial communities analyzed. B) NMDS ordination based on UniFrac distances of the microbial communities analyzed. In both ordinations, community type (single or pairwise mixture) is represented by different shapes, while each component (individual communities or their mixture) is represented by a different color. A small amount of jitter was added to avoid point overlap. C) Boxplot showing Bray–Curtis distances between samples from individual communities (intra-group) and between samples from the coalesced community and each of the individual communities (inter-group). The upper and lower box boundaries represent the 75th and 25th percentiles, respectively, and the line inside each box indicates the median. Whiskers represent the range of the data excluding outliers, defined as values located more than 1.5 times the interquartile range below the first quartile or above the third quartile. D) Boxplot showing UniFrac distances between samples from individual communities (intra-group) and between samples from the coalesced community and each of the individual communities (inter-group). Boxplot elements are defined as in panel C. E) Euler diagram quantifying the ASVs contributed by each community. Circle size is proportional to the number of ASVs present in each community, and shared ASVs are represented by overlapping regions. F) Rainbow plot based on ASV relative abundances. Each point represents one ASV, positioned along the y-axis according to its relative abundance. The x-axis represents each community type, including the two individual communities and their coalesced mixture. Different colors indicate ASV distribution patterns according to the communities in which they are detected (present in one individual community and the mixture, in both individual communities, in both individual communities and the mixture, or only in one community). A small amount of jitter was added to avoid point overlap.

**Figure 3.**
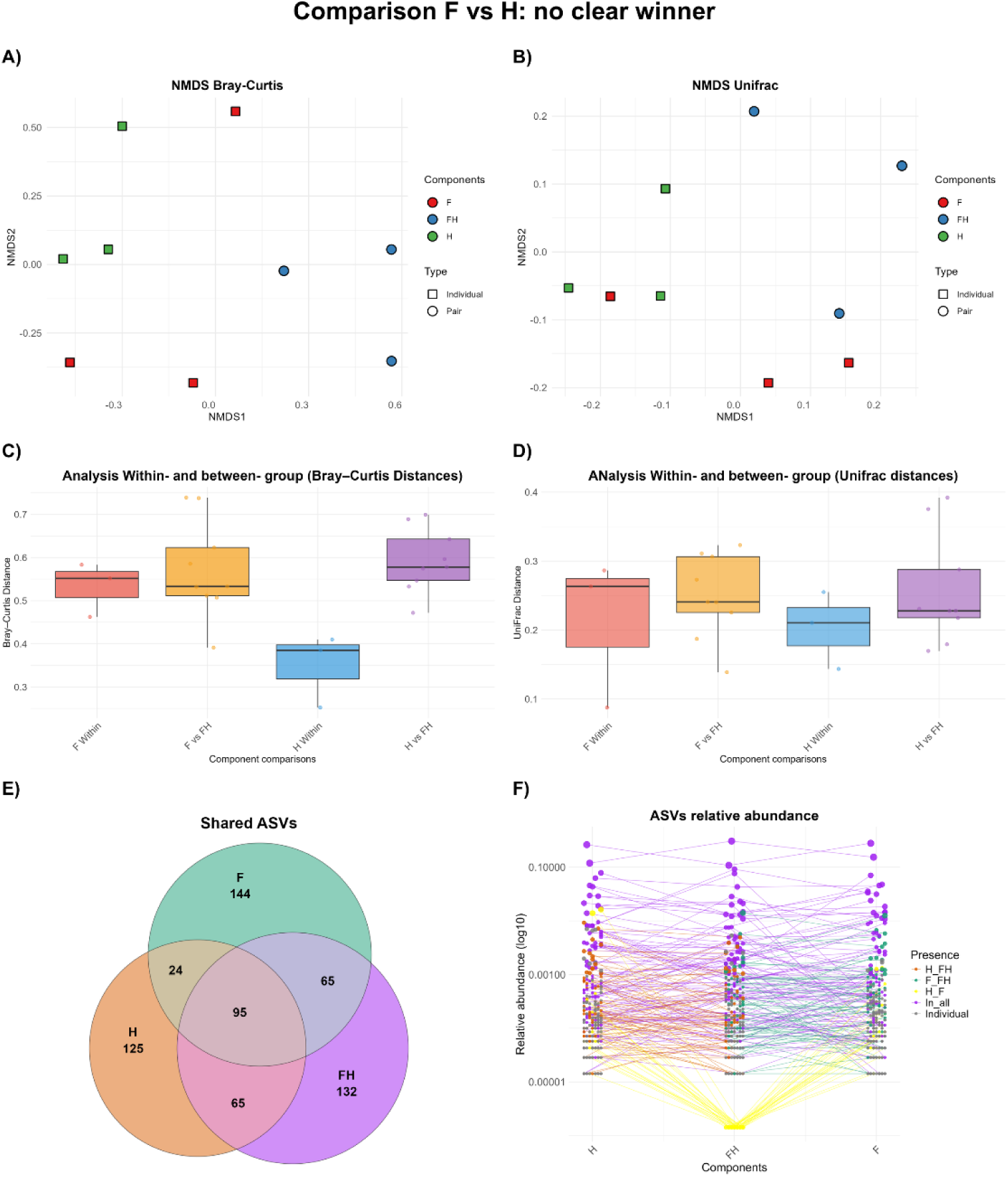
Representation of the different analyses performed to characterize the microbial communities resulting from community coalescence (FvsH). A) NMDS ordination based on Bray– Curtis distances of the microbial communities analyzed. B) NMDS ordination based on UniFrac distances of the microbial communities analyzed. In both ordinations, community type (single or pairwise mixture) is represented by different shapes, while each component (individual communities or their mixture) is represented by a different color. A small amount of jitter was added to avoid point overlap. C) Boxplot showing Bray–Curtis distances between samples from individual communities (intra-group) and between samples from the coalesced community and each of the individual communities (inter-group). The upper and lower box boundaries represent the 75th and 25th percentiles, respectively, and the line inside each box indicates the median. Whiskers represent the range of the data excluding outliers, defined as values located more than 1.5 times the interquartile range below the first quartile or above the third quartile. D) Boxplot showing UniFrac distances between samples from individual communities (intra-group) and between samples from the coalesced community and each of the individual communities (inter-group). Boxplot elements are defined as in panel C. E) Euler diagram quantifying the ASVs contributed by each community. Circle size is proportional to the number of ASVs present in each community, and shared ASVs are represented by overlapping regions. F) Rainbow plot based on ASV relative abundances. Each point represents one ASV, positioned along the y-axis according to its relative abundance. The x-axis represents each community type, including the two individual communities and their coalesced mixture. Different colors indicate ASV distribution patterns according to the communities in which they are detected (present in one individual community and the mixture, in both individual communities, in both individual communities and the mixture, or only in one community). A small amount of jitter was added to avoid point overlap.

**Figure 4.**
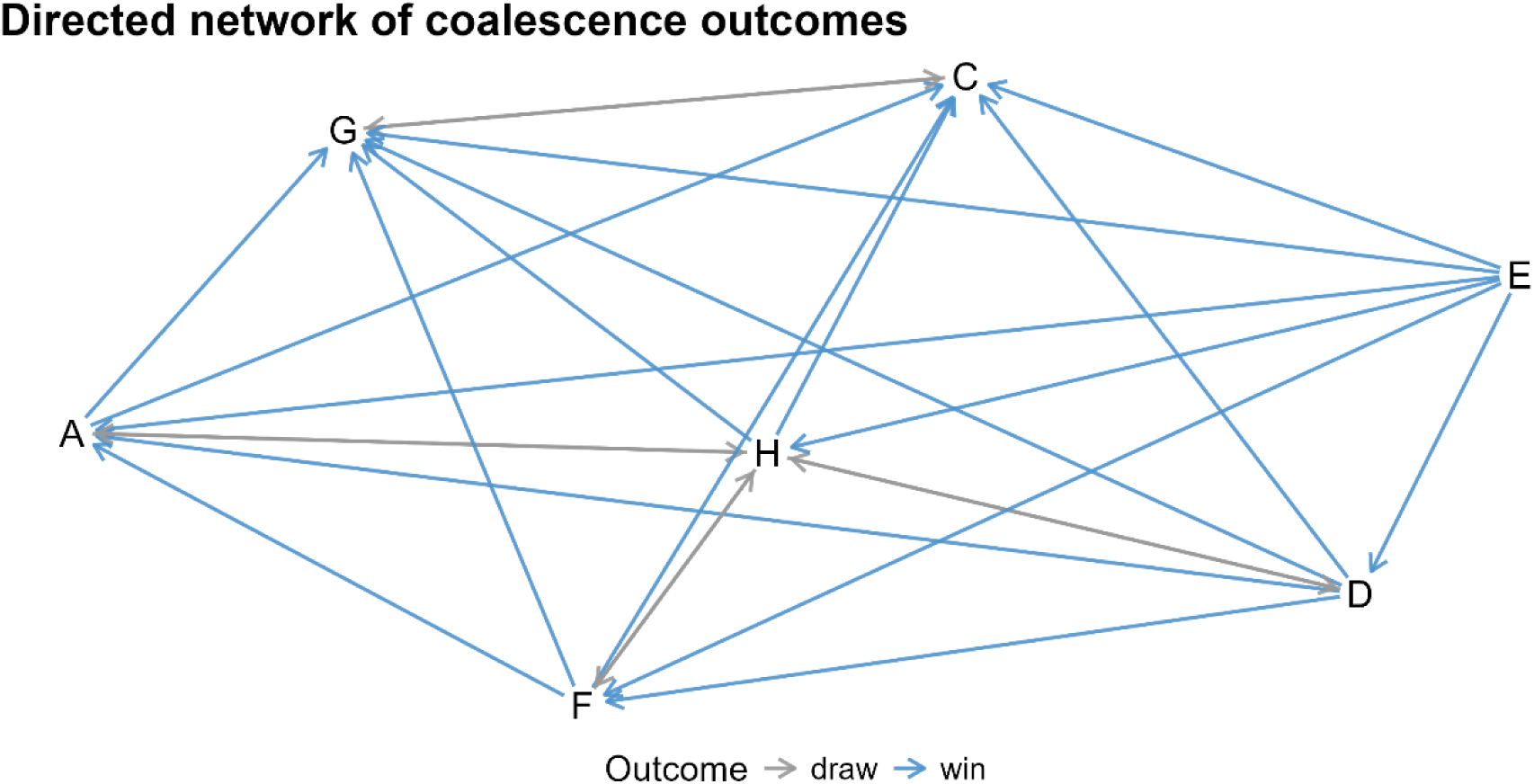
Directed network of coalescence outcomes. Dominance of one community over another is represented by a blue arrow (arrow points from dominant to dominated), whereas cases in which no clear winner emerged are represented by a grey line.

The effects of ASV richness (hereafter, Richness) and phylogenetic diversity (Faith’s index; hereafter, Diversity) on bacterial load (hereafter, Load) and plant dry weight (hereafter, Weight) were analyzed. Since Richness and Diversity were highly correlated (r = 0.85; P < 0.01), the analysis was performed using both Richness and the fraction of Diversity independent of Richness. To assess these effects, two simple linear models were fitted, and the significance of the effects was evaluated using Type II ANOVA. In the model fitted for Load, Richness did not exert a significant effect (P = 0.2). By contrast, the fraction of Diversity independent of Richness showed a weak effect on Load (F₁,₁₆₁ = 3.00; p = 0.08), with a marginal positive coefficient (β = 2.68 × 10⁻⁴; SE = 1.54 × 10⁻⁴; t = 1.73; P = 0.08), indicating a tendency for higher values of the Diversity fraction independent of Richness to be associated with increased Load. When evaluating the effects of these variables on Weight, Richness showed a weak overall effect (F₁,₁₆₁ = 2.85; P = 0.09), with a marginal positive coefficient (β = 3.38 × 10⁻⁵; SE = 2.00 × 10⁻⁵; t = 1.68; P = 0.09), indicating a tendency for increased Richness to be associated with greater Weight. In contrast, the fraction of Diversity independent of Richness did not significantly affect Weight (P = 0.6).

Three ASVs showed significant positive associations with Weight: ASV_117 (ρ = 0.459, FDR = 0.023), ASV_106 (ρ = 0.458, FDR = 0.023), and ASV_20 (ρ = 0.293, FDR = 0.037). These ASVs were taxonomically affiliated with Rhizobiaceae/*Ensifer*, Micrococcaceae/*Arthrobacter*, and Micrococcaceae/*Arthrobacter*, respectively. No ASVs showed significant negative associations with Weight. After correcting Weight for the mean effect of inoculum combination, the same ASVs remained among the strongest positive candidates, but none remained significant after multiple-testing correction. At broader taxonomic resolution, significant positive associations with Weight were also detected. At the genus level, *Mycobacterium* (ρ = 0.310, FDR = 0.005) and *Agromyces* (ρ = 0.260, FDR = 0.038) showed significant positive associations. At the family level, Mycobacteriaceae (ρ = 0.304, FDR = 0.003), Nocardiaceae (ρ = 0.270, FDR = 0.009), and Xanthomonadaceae (ρ = 0.246, FDR = 0.019) were significantly positively associated with Weight. All genus- and family-level associations lost significance after correcting for inoculum- combination effects.

No significant association was detected between Weight and distance to the global compositional centroid, either in the raw analysis (ρ = 0.065, P = 0.409) or after correcting for inoculum composition (ρ = 0.023, P = 0.769). To further investigate candidate growth-associated taxa, ASV_106 and ASV_117 were analyzed jointly. Co-occurrence of both ASVs was rare, occurring in only six samples across multiple inoculum combinations. Samples containing both ASVs showed substantially higher Weight than the rest of the dataset (mean = 0.043 g vs 0.028 g), and this difference was significant in a Wilcoxon rank-sum test (P = 0.0035).

We examined global community structure using Bray–Curtis NMDS ordinations. When considering single-inoculum communities only (Fig. 5A), samples clustered strongly according to inoculum identity, indicating that initial inoculum composition exerted a strong effect on rhizosphere community assembly. However, when all rhizosphere communities derived from single, pair, and triplet inocula were analyzed together (Fig. 5B), samples formed a largely continuous cloud with no clear evidence of discrete compositional clusters or alternative community states.

**Figure 5.**
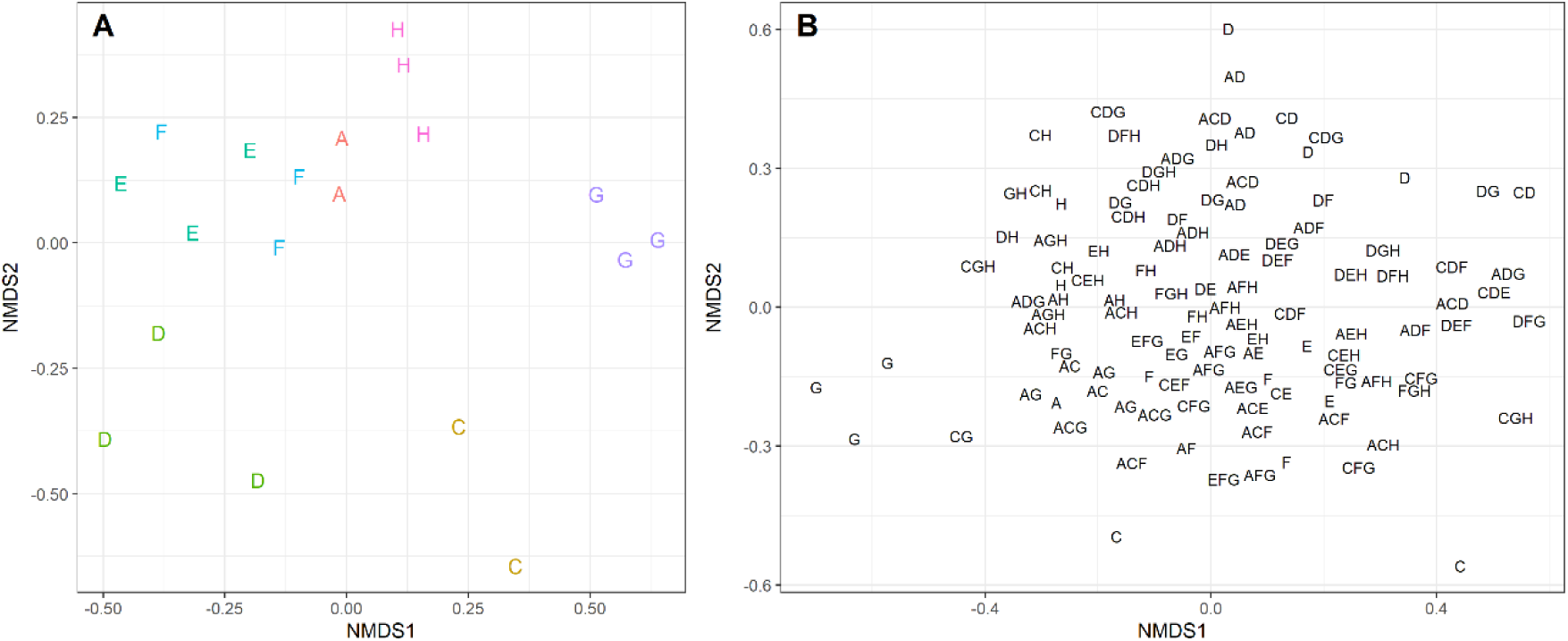
Non-metric multidimensional scaling (NMDS) ordination based on Bray–Curtis dissimilarities showing the compositional relationships among rhizosphere microbial communities across inoculum contexts. Each point represents an individual sample, and distances between points reflect differences in community composition, with shorter distances indicating more similar communities. **(A)** NMDS including only rhizosphere communities derived from single inocula. Samples are colored according to inoculum identity (A, C, D, E, F, G, and H). **(B)** NMDS including all samples from single, pairwise, and triplet inoculum combinations. Samples are labeled according to their inoculum composition, where single letters indicate single inocula, two-letter labels indicate pairwise combinations, and three-letter labels indicate triplet combinations.

We next analyzed rank–abundance distributions to determine whether increasing inoculum complexity altered the internal structure of assembled communities (Fig. 6). Surprisingly, single, pair, and triplet-derived communities exhibited highly similar rank–abundance profiles, with little evidence of systematic differences in dominance structure across inoculum complexities. Across all samples, only two ASVs were typically sufficient to explain approximately 50% of total relative abundance (median = 2; IQR = 2–3), whereas a median of 19 ASVs explained 90% of total abundance (IQR = 12–28). This pattern was strongly associated with dominance by *Pseudomonas*. The two most abundant ASVs were both assigned to Pseudomonas in 83.5% of samples, and in most remaining samples *Pseudomonas*-affiliated ASVs still occupied the first-ranked position. Although 186 ASVs in the dataset were taxonomically assigned to *Pseudomonas*, dominance was concentrated in only two ASVs, ASV_1 and ASV_2, which accounted for 29.3% and 19.1% of total relative abundance (Supplementary Figure 22), respectively. Only four samples lacked *Pseudomonas* among their two most abundant taxa, and all belonged to G-containing inocula (G, CG, or ADG). These alternative states were characterized by ASVs affiliated with *Burkholderia* and/or *Arthrobacter*, corresponding to ASV_10 and ASV_9. This pattern was consistent with the source distribution of the dominant ASVs; *Pseudomonas* ASV_1 was not detected in the G single inoculum, whereas the alternative dominant ASVs ASV_9 and ASV_10 were detected among single inocula mostly in G (Supplementary figure 23). Moreover, these dominant configurations were associated with different accompanying taxa: communities dominated by *Pseudomonas* ASV_1 or ASV_2 were more frequently accompanied by *Chryseobacterium* and *Flavobacterium*, whereas communities dominated by ASV_9 or ASV_10 were more frequently associated with *Acidovorax* and *Xanthomonas*. However, G-containing communities did not consistently follow this alternative trajectory, as some replicates converged to the canonical *Pseudomonas*-dominated configuration whereas others shifted toward the alternative dominant states.

**Figure 6.**
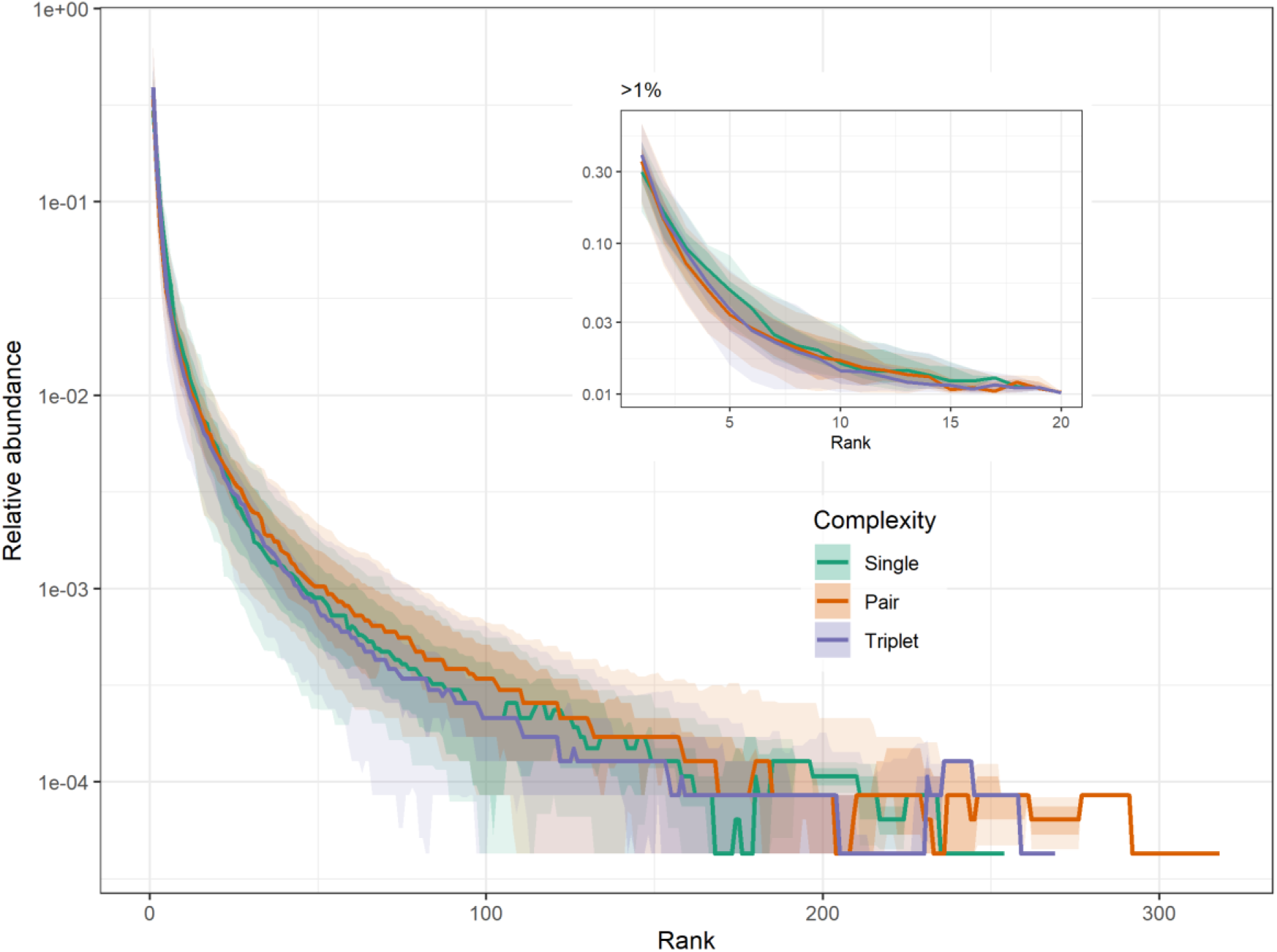
Rank–abundance distributions of rhizosphere microbial communities across inoculum complexity levels. Curves show the median rank–abundance profiles for communities derived from single, pairwise, and triplet inoculum combinations, with shaded bands representing the 25–75% and 5–95% percentile ranges across samples. Relative abundance is shown on a logarithmic scale, and ASVs are ranked within each sample from most to least abundant. The inset displays the same analysis restricted to ASVs with relative abundance >1.

We assessed reproducibility by comparing Pearson correlations between samples belonging to the same inoculum combination and samples belonging to different combinations (Fig. 7). Intra- combination correlations were higher than inter-combination correlations, confirming that inoculum identity influenced final community composition. However, intra-combination correlations were similar among single, pair, and triplet-derived communities, suggesting that increasing inoculum complexity did not produce a detectable increase in replicate-to-replicate stochasticity at the level of whole-community structure.

**Figure 7.**
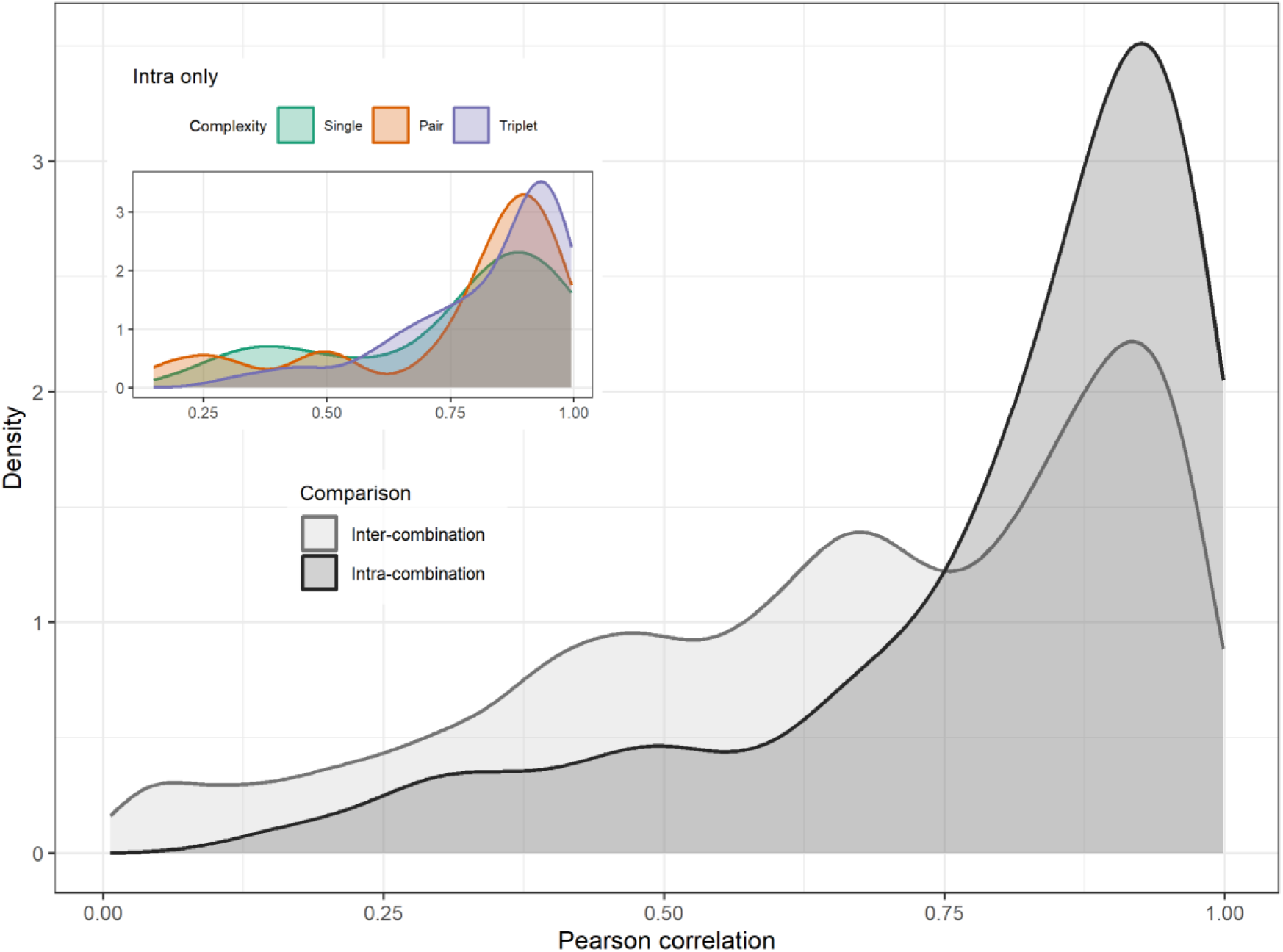
Distribution of pairwise Pearson correlations between rhizosphere community compositions across samples. The main panel compares the density distributions of correlations between samples originating from the same inoculum combination (intra-combination) and from different inoculum combinations (inter-combination). The inset shows the intra-combination correlation distributions stratified by inoculum complexity.

To quantify convergence toward a common community state, we defined a global rhizosphere compositional centroid as the mean relative abundance profile across all samples and calculated Bray–Curtis distances from each sample to this centroid. Triplet-derived communities showed a narrower distribution of distances and were slightly more concentrated around the centroid than pair-derived communities, suggesting greater convergence toward the average rhizosphere composition with increasing inoculum complexity (Fig. 8). In contrast, pair-derived communities displayed substantially broader distance distributions, indicating greater compositional heterogeneity. Single inocula occupied distinct positions relative to the centroid. Communities derived from D, E, and F were closest to the global centroid, indicating that these inocula already resembled the dominant rhizosphere configuration. In contrast, A and C were located at intermediate distances, whereas G was by far the most distant single inoculum.

**Figure 8.**
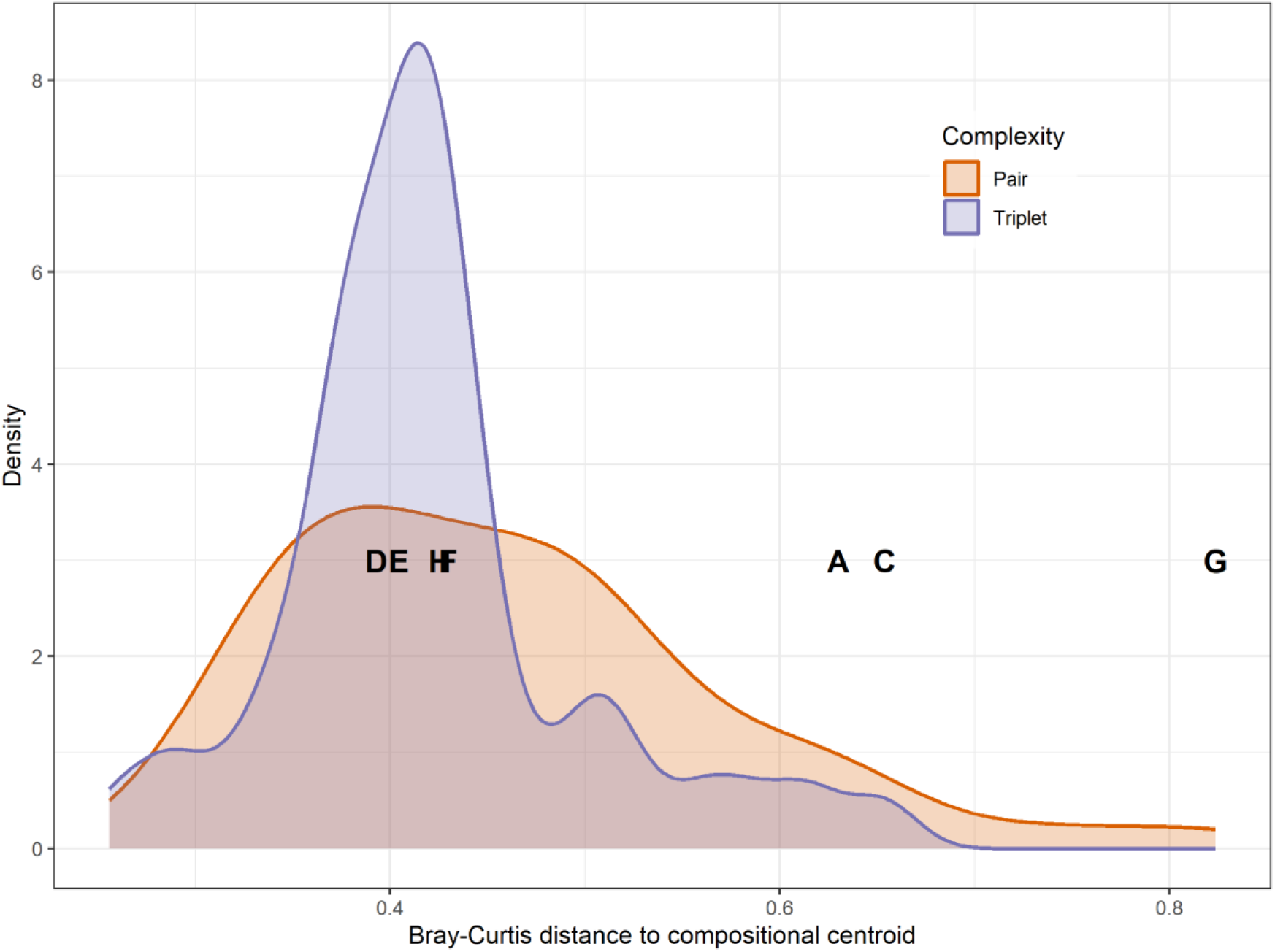
Bray–Curtis distance of rhizosphere communities to the global compositional centroid. The global centroid was defined as the mean relative abundance profile across all samples and represents the average rhizosphere community composition. Density curves show the distribution of Bray–Curtis distances to the centroid for communities derived from pairwise and triplet inoculum combinations. Single-inoculum communities are represented by letters positioned at their mean distance to the centroid.

Finally, ASV conditional persistence and mean retention were positively associated (Fig. 9 Left), indicating that ASVs maintaining higher abundance after mixing were also more likely to persist across community contexts. Most ASVs occupied the lower-left region of the trait space, characterized by strongly negative retention values and low persistence, indicating that the majority of taxa detected in single inocula were strongly reduced or lost following community mixing. By contrast, a relatively small number of ASVs occupied the upper-right region, characterized by high persistence and comparatively high retention, consistent with strong competitive performance across multiple assembly contexts. ASVs with broader source distributions tended to show higher persistence and retention. In the second trait-space projection (Fig. 9 Right), context dependence was highest among ASVs with intermediate-to-low retention, whereas ASVs with the highest retention values generally exhibited low context dependence, indicating that the most successful colonizers were also the most robust to changes in community context.

**Figure 9.**
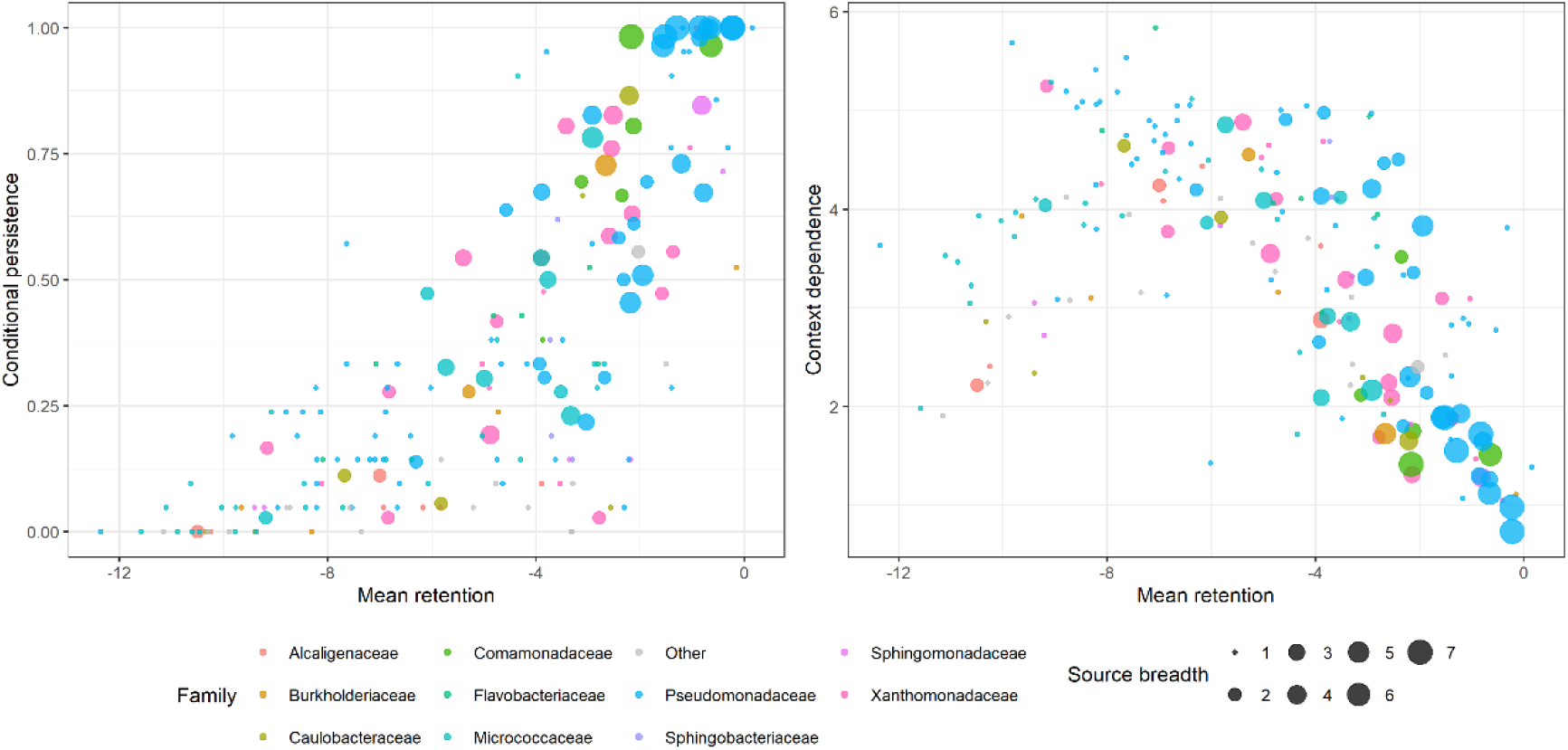
Relationships between ASV retention and two derived assembly metrics across rhizosphere communities. Each point represents one ASV. In both panels, the x-axis shows mean retention across samples. **Left panel:** conditional persistence as a function of mean retention. **Right panel:** context dependence as a function of mean retention. Points are colored according to taxonomic family, with major families shown individually and all remaining families grouped as “Other”. Point size indicates source breadth, defined as the number of inoculum sources in which each ASV was detected.

## DISCUSSION

Despite minor imbalance in the experimental design caused by the loss of a small number of replicates due to failed plant growth or insufficient sequencing depth, the dataset retained high replication across inoculum combinations and provided strong resolution to investigate rhizosphere community assembly across a broad range of assembly contexts. The combination of replication, multiple inoculum compositions, and controlled coalescence events provided a robust framework to explore how initial community composition, microbial interactions, and host filtering jointly shape early rhizosphere microbiome assembly in tomato.

Notwithstanding the large compositional differences among the original soil inocula ^30^, rhizosphere assembly was strongly constrained by host filtering, as evidenced by the highly similar family-level composition observed across most assembled communities. At the same time, the composition of the initial inoculum remained an important determinant of community assembly, as single-inoculum rhizosphere communities clustered strongly according to inoculum identity. As previously reported ^11,30,31^, these results support the view that rhizosphere assembly emerges from the interaction between strong plant-driven ecological filtering and the composition of the source microbial pool.

However, when communities derived from single, pair, and triplet inocula were analyzed jointly, no clear evidence of discrete compositional clusters or stable alternative community states emerged. Instead, samples formed a largely continuous compositional cloud, suggesting that different assembly trajectories generally converged within the same broad ecological landscape rather than toward sharply separated attractors.

This overall convergence was further reflected in the strong and recurrent dominance of *Pseudomonas*-affiliated ASVs. Although 186 ASVs were taxonomically assigned to *Pseudomonas*, dominance was overwhelmingly concentrated in only two ASVs, which occupied the highest abundance ranks in most communities. This pattern suggests that rhizosphere filtering operated not only at broad taxonomic levels but also at much finer ecological resolution, strongly favoring a small subset of highly competitive colonizers.

Notably, limited evidence for an alternative assembly trajectory was observed in communities containing inoculum G. In these cases, *Pseudomonas* dominance was occasionally replaced by ASVs affiliated with *Burkholderia* and *Arthrobacter*. However, this alternative state was not consistently reproduced, as G-containing communities showed heterogeneous outcomes: some replicates converged toward the canonical *Pseudomonas*-dominated configuration, whereas others shifted toward the alternative dominant state. This suggests that inoculum G increased the probability of accessing an alternative assembly trajectory without imposing a fully stable alternative community state.

These observations are also consistent with a lottery-like assembly process ^32^, in which multiple taxa with partially overlapping ecological functions compete for similar niches, but only one successfully establishes dominance. Under this framework, *Pseudomonas* would represent the most frequent winner of a dominant early-colonizer niche, whereas *Burkholderia* or *Arthrobacter* may occasionally occupy similar functional space under specific source-community contexts, particularly in G-derived inocula.

One of the most striking findings of this study was the remarkable conservation of overall community architecture across assembly contexts. Increasing inoculum complexity from single to pairwise and triplet combinations did not produce a detectable increase in replicate-to-replicate stochasticity at the whole-community level, as intra-combination correlations remained broadly similar across all inoculum classes.

Even more strikingly, rank–abundance distributions were highly similar across single-, pair-, and triplet-derived communities. Despite large differences in initial inoculum composition, richness, and coalescence complexity, rhizosphere communities consistently converged toward a highly uneven abundance structure dominated by a small number of highly abundant ASVs and a long tail of low-abundance taxa. In most communities, only two ASVs were sufficient to explain approximately half of the total relative abundance, whereas a median of just 19 ASVs explained 90% of total abundance.

These results suggest that the strongest deterministic feature of rhizosphere assembly in this system may not be convergence toward identical taxonomic composition, but rather convergence toward a highly reproducible structural organization of the community. In other words, the tomato rhizosphere appears to constrain not only which taxa can persist, but also the overall abundance hierarchy that emerges during assembly.

This pattern is consistent with macroecological observations showing that microbial communities frequently display strongly skewed abundance distributions composed of a few highly successful taxa and a long rare biosphere ^33^. It also closely resembles the “emergent simplicity” described by Goldford et al. (2018) ^34^, where taxonomically distinct microbial communities assembled under common environmental conditions converged toward recurrent community-level structures. More broadly, these observations support the idea that relatively simple ecological constraints may govern the assembly of complex microbial communities, producing predictable structural outcomes despite substantial variation in initial conditions ^35^.

To further investigate whether the recurrent community architecture observed across assembly contexts reflected convergence toward a common rhizosphere state, we quantified the distance of each community to a global rhizosphere compositional centroid. This analysis further supported strong deterministic assembly, as increasing inoculum complexity promoted convergence toward the centroid: triplet-derived communities were more tightly clustered around the centroid than pair-derived communities, whereas pair-derived communities showed greater heterogeneity. Thus, rather than increasing stochasticity, mixing a larger number of source communities reduced compositional variability and drove stronger convergence toward a common rhizosphere configuration.

The positions of single inocula relative to the centroid were particularly informative. Communities derived from D, E, and F were closest to the centroid, suggesting that these inocula already resembled the dominant rhizosphere configuration, whereas A and C occupied intermediate positions and G was by far the most distant inoculum. This pattern closely mirrored the dominance hierarchy observed in the coalescence experiments. Communities D and E consistently showed the highest dominance rates, whereas C and especially G were among the weakest competitors. Thus, proximity to the rhizosphere centroid appeared to be a strong predictor of competitive success during coalescence. Because the global centroid was calculated from the full dataset, including single-, pair-, and triplet-derived communities, its position is necessarily influenced by the composition of the communities analyzed. Thus, the observed association between centroid proximity and coalescence dominance should be interpreted as a descriptive relationship rather than as fully independent evidence that proximity to a predefined rhizosphere attractor determines competitive success.

This suggests that communities already closer to the canonical rhizosphere state may possess a greater proportion of taxa pre-adapted to the selective conditions imposed by the tomato rhizosphere, giving them a competitive advantage during coalescence. Consistent with this interpretation, the strongest competitors (D and E) originated from orchard garden soils, whereas weaker competitors such as C and G originated from forest and dryland agricultural soils. Although speculative, this raises the possibility that historical contingency and prior exposure to tomato or other horticultural crop rhizospheres may have pre-conditioned certain communities for more efficient colonization of the tomato rhizosphere. G represented the clearest exception to the canonical rhizosphere attractor. Its large distance from the centroid was consistent with its role as the main source of alternative assembly trajectories, suggesting that G contained community configurations with the greatest potential to deviate from the dominant rhizosphere state. However, even G did not consistently escape the canonical attractor, as some G-containing communities converged toward the dominant rhizosphere configuration while others followed alternative trajectories.

With respect to host phenotype, we found only weak associations between broad community descriptors and plant performance; Richness showed a marginal positive association with plant biomass, whereas Diversity independent of richness showed a marginal positive association with bacterial load.

Several microbial taxa showed significant positive associations with plant biomass. However, most of these associations disappeared after correcting for inoculum combination effects, indicating that their apparent effects were strongly influenced by community context. This indicates that the apparent taxon-level associations were strongly confounded by inoculum- combination context and should not be interpreted as independent effects of individual taxa. More broadly, these results support the idea that plant performance may reflect properties of the broader assembled community rather than the independent effects of single taxa and caution against interpreting taxon-level correlations with plant phenotype as evidence of direct causal effects.

## Conclusion

Overall, the replication and broad range of inoculum combinations in this controlled host-associated experimental system allowed us to characterize rhizosphere microbiome assembly across an unusually wide range of assembly contexts. The most striking result was the recurrent emergence of a highly conserved community architecture across all treatments. Despite major differences in inoculum composition and richness, rhizosphere communities consistently converged toward the same strongly uneven abundance structure dominated by a small number of taxa.

Increasing inoculum complexity drove communities progressively closer to a common rhizosphere centroid, and coalescence outcomes, particularly community dominance hierarchies, closely mirrored the distance of the original inocula to this centroid. Together, these patterns are consistent with the existence of a canonical rhizosphere attractor in this system.

One possible explanation is that this simplified experimental environment, largely defined by root-derived resources in the absence of soil complexity, supports a limited number of dominant ecological niches. Under this framework, increasing inoculum complexity increases the probability that the most competitive colonizers for each niche are present, thereby promoting stronger convergence toward the canonical rhizosphere state. The exploratory association between source soil-use history and dominance further raises the possibility that historical contingency may contribute to coalescence outcomes, potentially reflecting prior adaptation to rhizospheres associated with horticultural cropping systems.

## Funding

This work was funded by the Spanish Ministry of Science and Innovation grant CNS2022-135371 awarded to DAC.

## Notes

https://github.com/microenvgen/Coalescence

### Competing Interest Statement

The authors have declared no competing interest.

### Summary of Updates

Several minor changes in wording and explanations.

