## Supplementary for "Combinatorial community coalescence in early tomato assembly reveals a rhizosphere attractor in composition and abundance architecture"

^1^Microbial and Environmental Genomics Group, Departamento de Biología, Universidad Autónoma de Madrid, 28049, Spain.

| **Soil name** | **A** | **C** | **D** | **E** | **F** | **G** | **H** |
| --- | --- | --- | --- | --- | --- | --- | --- |
| **Origin** | Guadalajara (Guadalajara) | Valdelatas (Madrid) | Vigo  (Pontevedra) | Ruiseñada (Cantabria) | Moraira  (Alicante) | Badajoz  (Badajoz) | Algete (Madrid) |
| **Soil Type** | recreational garden | Forest | recreational garden | recreational garden | ruderal soil | dryland cultivation soil | ruderal soil |
| **Texture Class** | Sandy  Clay  Loam | Loamy  Sand | Loamy  Sand | Loamy  Sand | Loamy  Clay | Loamy  Sand | Sandy  Clay  Loam |
| **Clay (%)** | 28 | 7 | 12 | 15 | 33 | 19 | 25 |
| **Sand (%)** | 14 | 10 | 21 | 24 | 26 | 24 | 10 |
| **Arena (%)** | 58 | 83 | 67 | 61 | 41 | 57 | 65 |
| **Conductivity**  **(μS/cm)** | 330 | 140 | 265 | 184 | 125 | 196 | 88,8 |
| **pH** | 7,46 | 5,84 | 5,46 | 6,8 | 7,74 | 6,42 | 7,02 |
| **Organic Matter**  **(%)** | 11,4 | 2,42 | 4,61 | 3,51 | 2,25 | 1,29 | 0,49 |
| **Available Nitrogen**  **(mg/kg)** | 5503 | 1112 | 2606 | 2,312 | 986 | 944 | 344 |
| **Available**  **Phosphorus**  **(mg/kg)** | 349 | 14,3 | 218 | 103 | < 9.80 | 35,4 | 13,8 |
| **Active Limestone**  **(%)** | 4,12 | < 0.500 | < 0.500 | < 0.500 | 7,81 | <0,5 | 1,13 |
| **Available Ca**  **(meq/100g)** | 21,1 | 3,84 | 6,86 | 11,3 | 14,9 | 6,05 | 8,99 |
| **Available Mg**  **(meq/100g)** | 4,6 | 1,07 | 0,9 | 1,12 | 0,88 | 2,21 | 1,57 |
| **Available**  **Potassium**  **(meq/100g)** | 3,46 | 0,41 | 0,94 | 0,61 | 0,39 | 0,37 | 0,26 |
| **Available Na**  **(meq/100g)** | 0,16 | < 0.05 | 0,1 | 0,08 | 0,09 | 0,25 | 0,06 |
| **C/N ratio** | 12 | 12,6 | 10,3 | 8,8 | 13,3 | 7,9 | 8,23 |
| **Latitude** | 40.64 | 40.54 | 42.20 | 43.36 | 38.69 | 38.85 | 40.61 |
| **Lonongitude** | -3.15 | -3.69 | -8.71 | -4.29 | 0.13 | -6.67 | -3.50 |
| **Elevation (m)** | 708 | 700 | 137 | 45 | 17 | 222 | 741 |

**Supplementary table 1. Characteristics of the soil samples used.**

| **Inoculum** | **Richness** | **Evenness** | **Diversity** | **Soil** |
| --- | --- | --- | --- | --- |
| **E** | 152 ± 25,9 | 2,37 ± 0,57 | 2230 ± 377 | Recreational orchad |
| **D** | 175 ± 80 | 2,57 ± 0,29 | 2710± 672 | Recreational orchad |
| **F** | 164 ± 39,7 | 2,77 ± 0,61 | 2650 ± 365 | Ruderal soil |
| **H** | 169 ± 61,1 | 3,03 ± 0,19 | 2810 ± 407 | Ruderal soil |
| **A** | 164 ± 24 | 3,32 ± 0,07 | 2650 ± 457 | Recreational orchad |
| **G** | 122 ± 20,7 | 2,51 ± 0,07 | 2650 ± 444 | Dryland cultivation soil |
| **C** | 85 ± 42,4 | 2,16 ± 0,20 | 2130 ± 577 | Forest |

**Supplementary table 2. Diversity metrics of the individual communities and the soil type from which each community originated.**

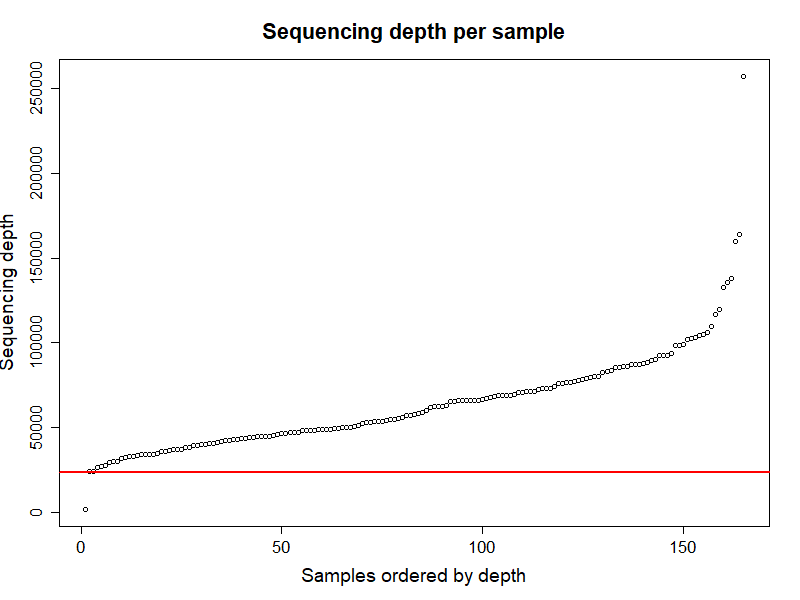

**Supplementary Figure 1. Sequencing depth per sample in the dataset.** Each point represents an individual sample (X-axis) ordered from lowest to highest sequencing depth and the corresponding number of reads per sample (Y-axis). The red line indicates the subsampling depth used (23,394 reads per sample).

**Supplementary Figures 2-21. Representation of the different analyses performed to characterize the microbial communities resulting from community coalescence. Each table’s title indicates the pair being analyzed.** A) NMDS ordination based on Bray–Curtis distances of the microbial communities analyzed. B) NMDS ordination based on UniFrac distances of the microbial communities analyzed. In both ordinations, community type (single or pairwise mixture) is represented by different shapes, while each component (individual communities or their mixture) is represented by a different color. A small amount of jitter was added to avoid point overlap. C) Boxplot showing Bray–Curtis distances between samples from individual communities (intra-group) and between samples from the coalesced community and each of the individual communities (inter-group). The upper and lower box boundaries represent the 75th and 25th percentiles, respectively, and the line inside each box indicates the median. Whiskers represent the range of the data excluding outliers, defined as values located more than 1.5 times the interquartile range below the first quartile or above the third quartile. D) Boxplot showing UniFrac distances between samples from individual communities (intra-group) and between samples from the coalesced community and each of the individual communities (inter-group). Boxplot elements are defined as in panel C. E) Euler diagram quantifying the ASVs contributed by each community. Circle size is proportional to the number of ASVs present in each community, and shared ASVs are represented by overlapping regions. F) Rainbow plot based on ASV relative abundances. Each point represents one ASV, positioned along the y-axis according to its relative abundance. The x-axis represents each community type, including the two individual communities and their coalesced mixture. Different colors indicate ASV distribution patterns according to the communities in which they are detected (present in one individual community and the mixture, in both individual communities, in both individual communities and the mixture, or only in one community). A small amount of jitter was added to avoid point overlap.

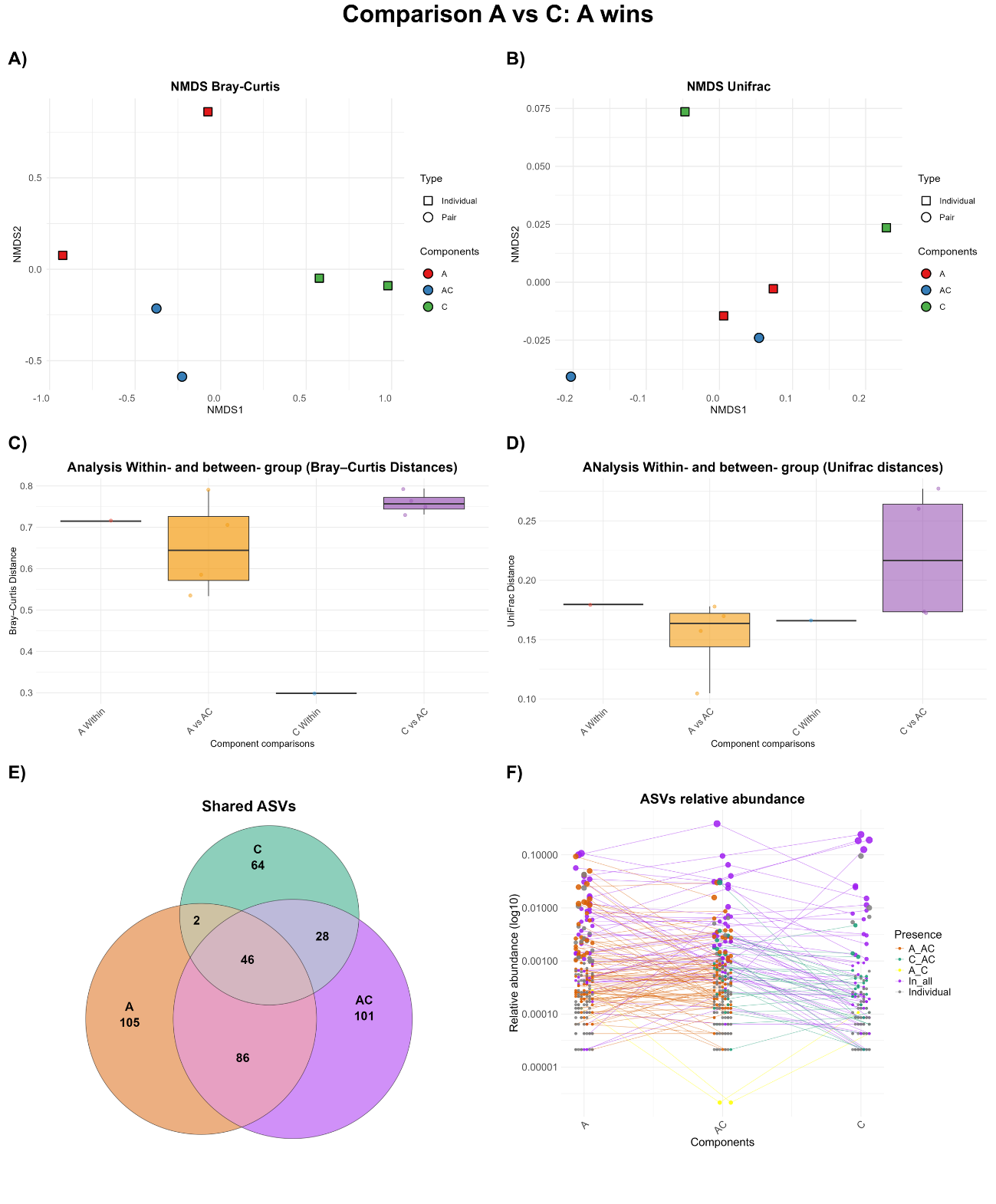

**Supplementary Figure 2**

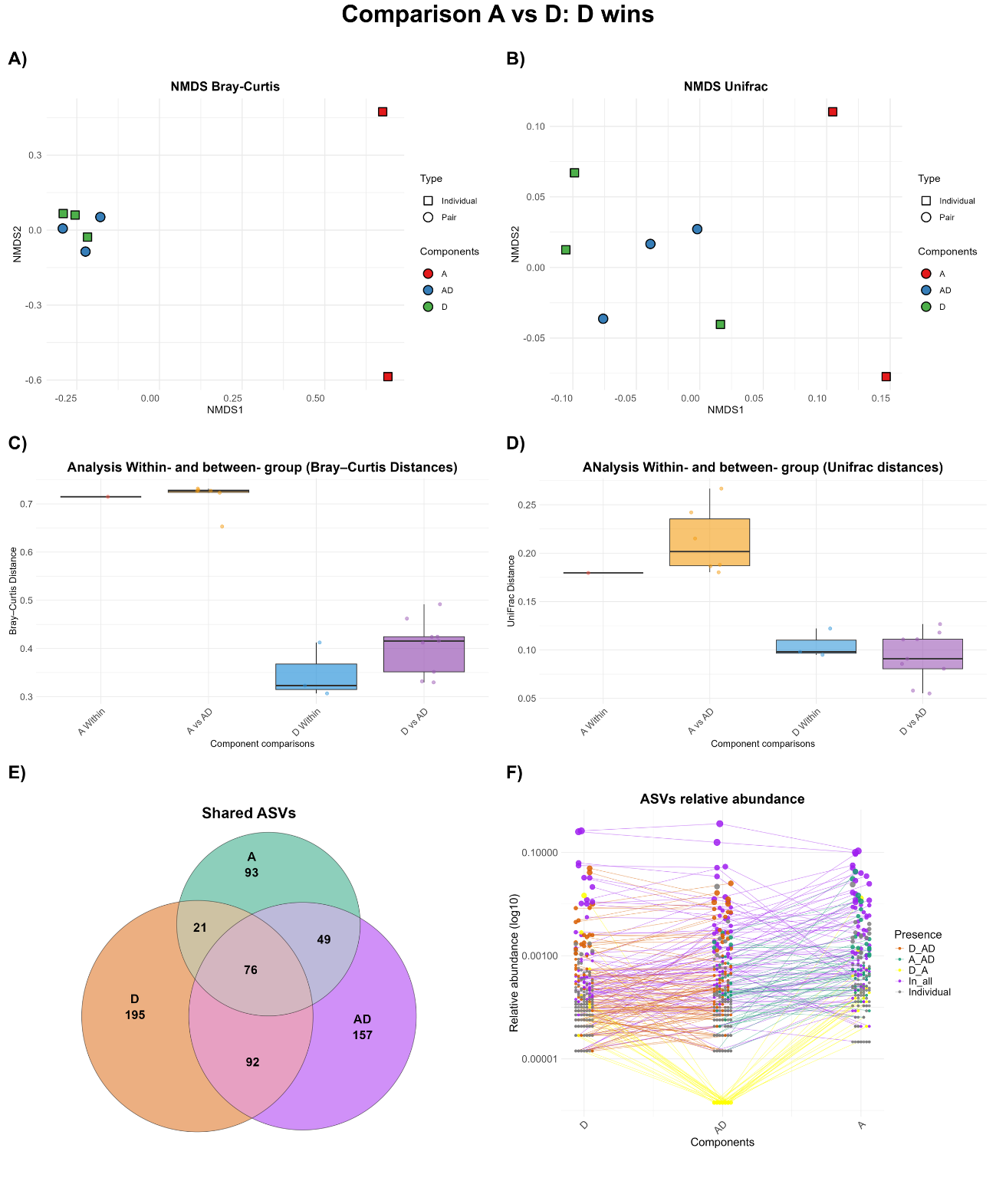

**Supplementary Figure 3**

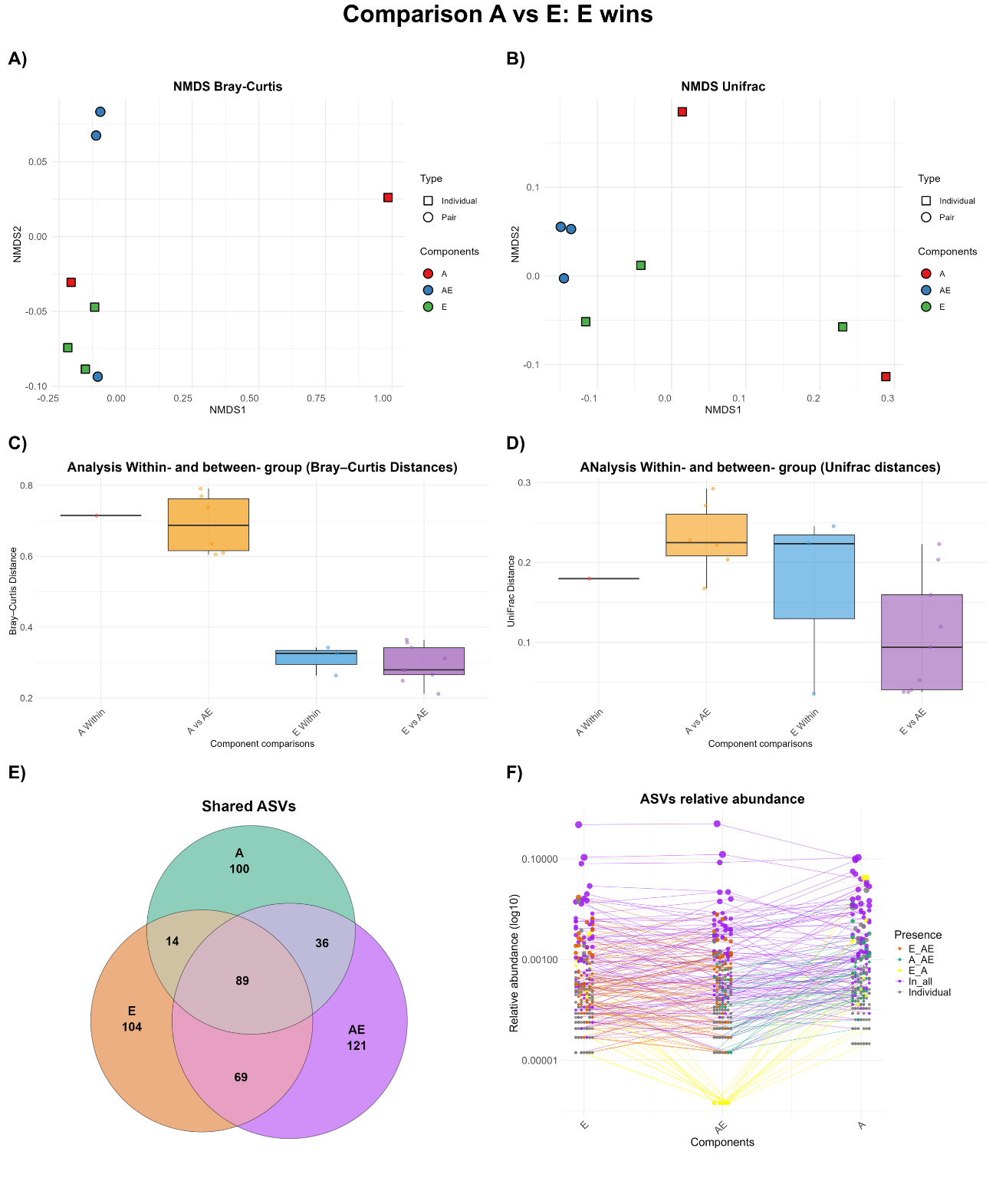

**Supplementary Figure 4**

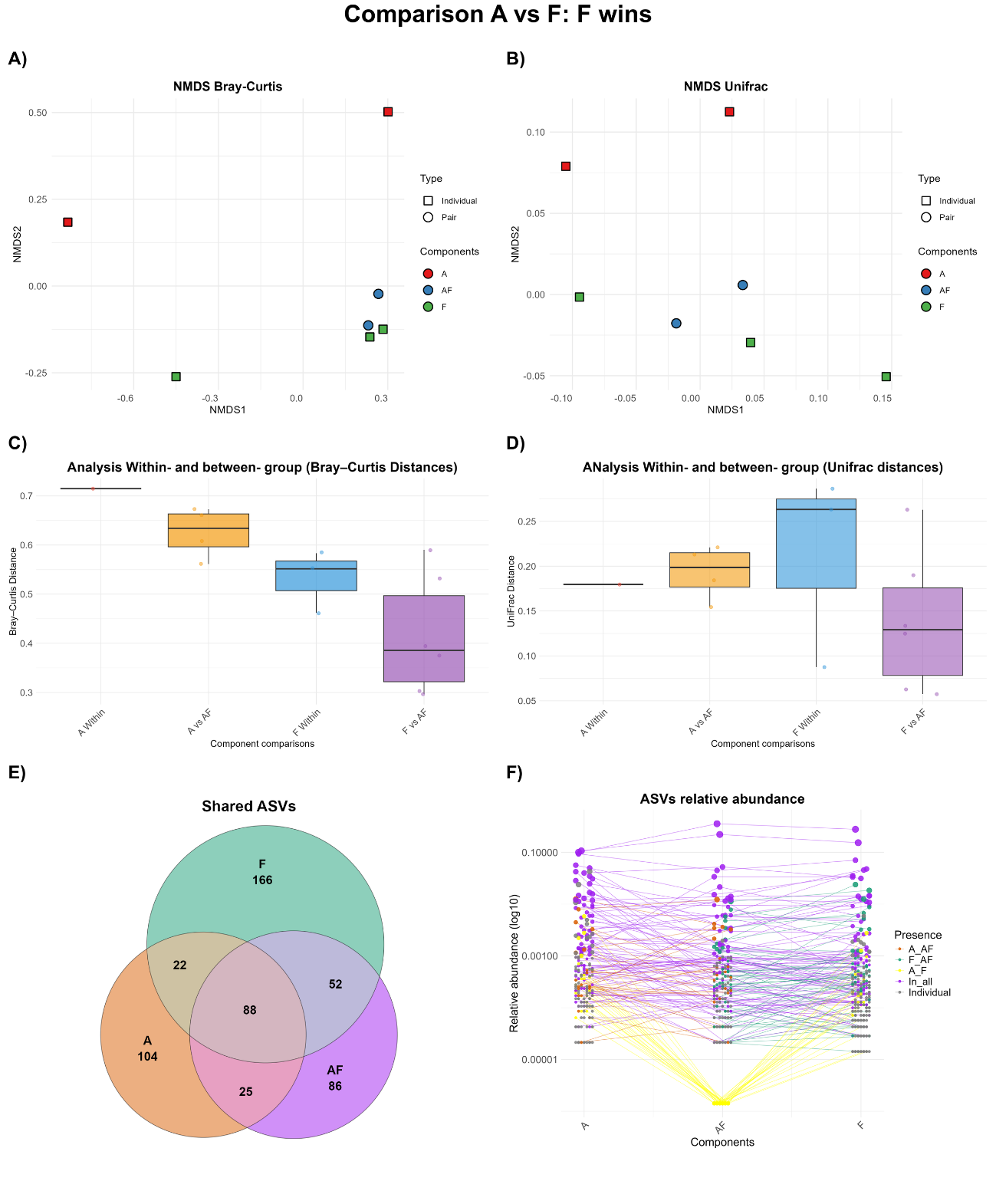

**Supplementary Figure 5**

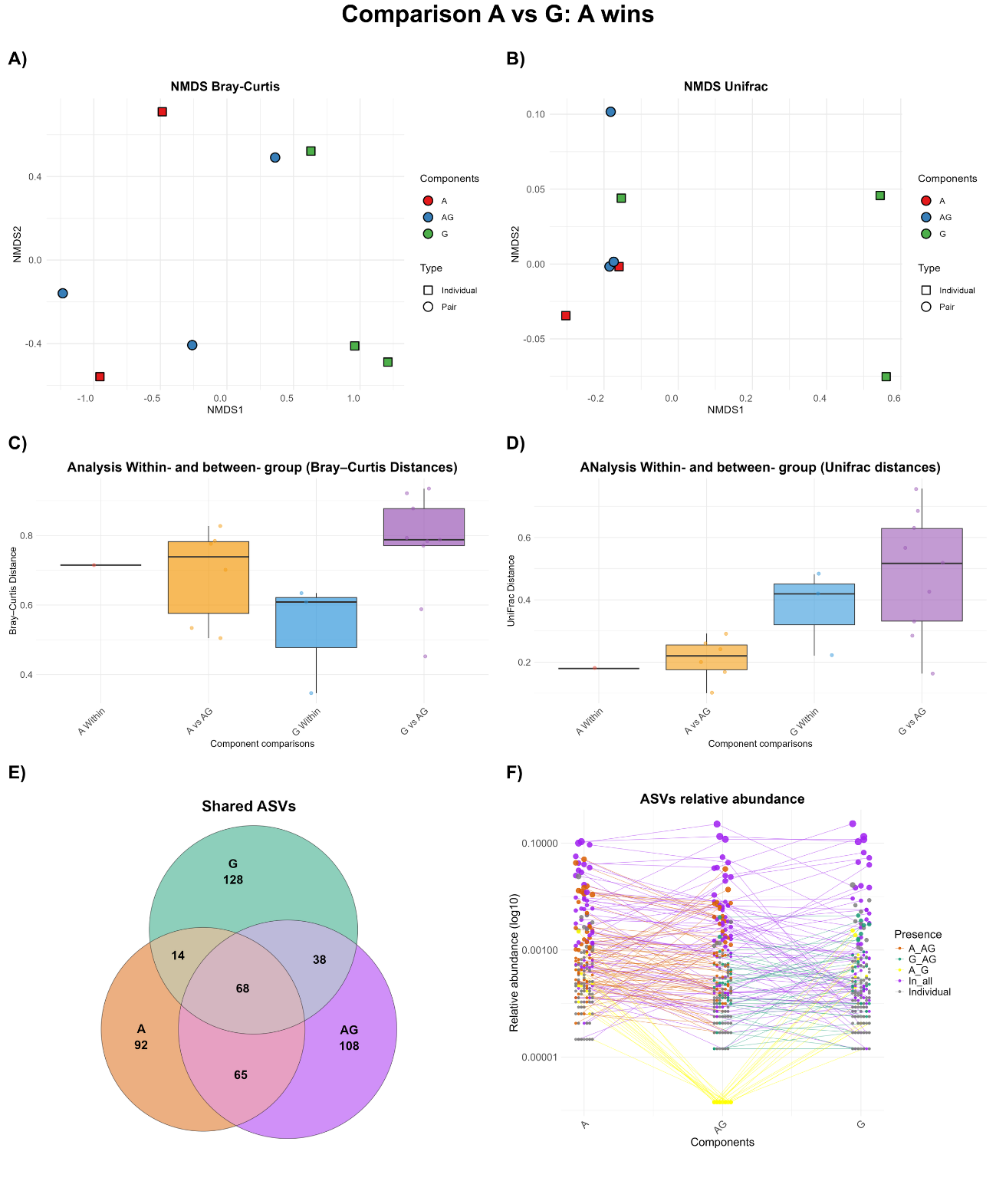

**Supplementary Figure 6**

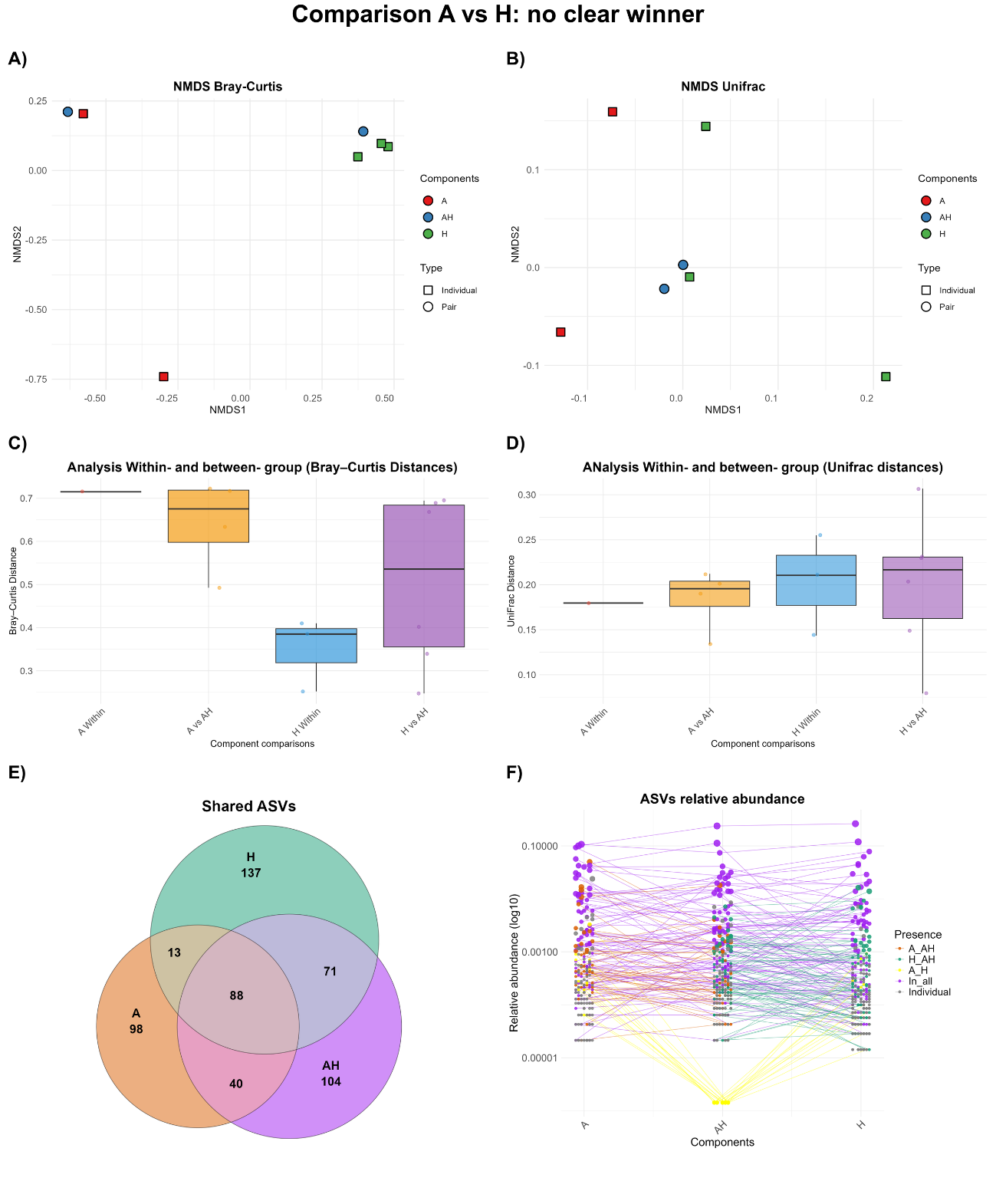

**Supplementary Figure 7**

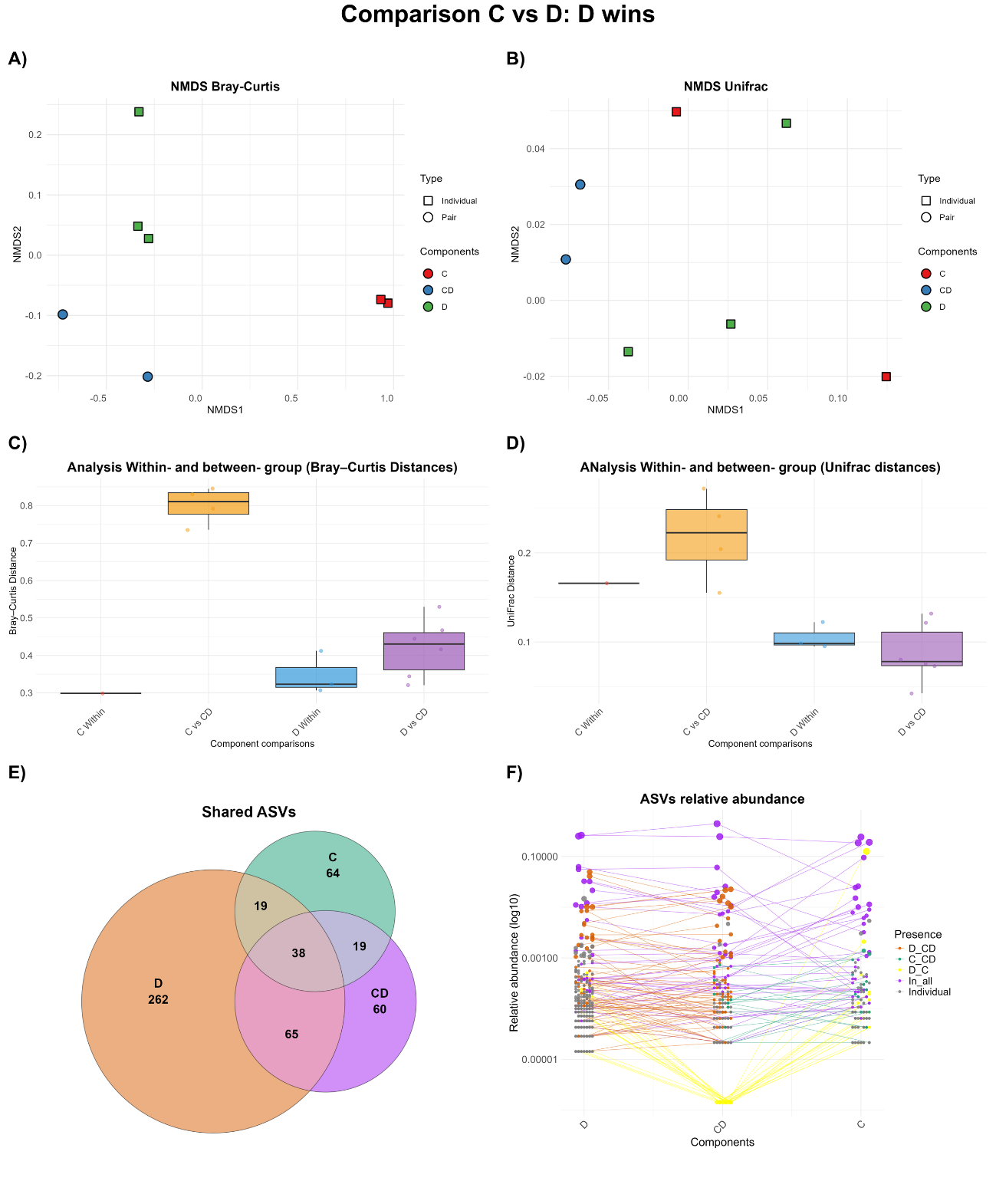

**Supplementary Figure 8**

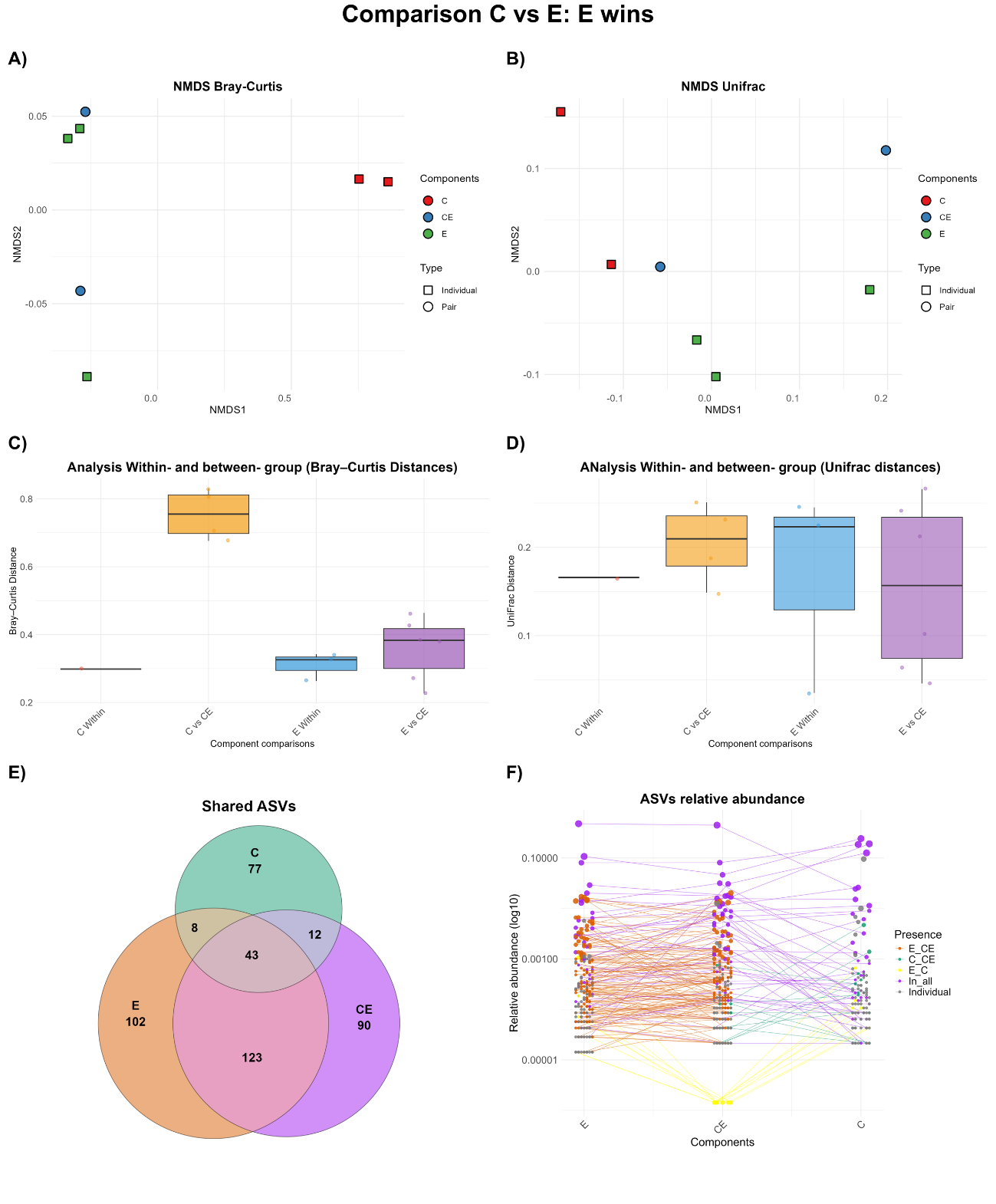

**Supplementary Figure 9**

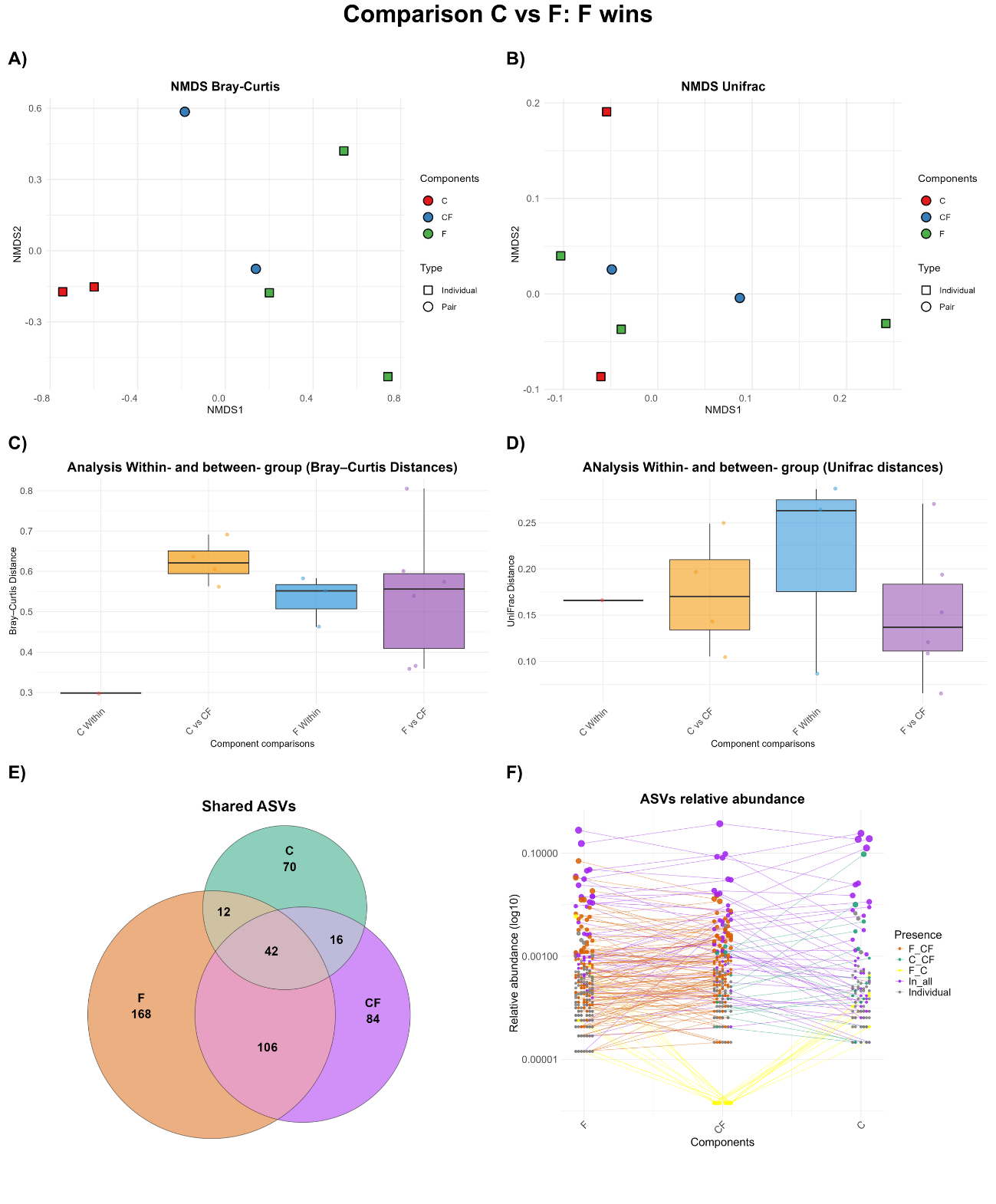

**Supplementary Figure 10**

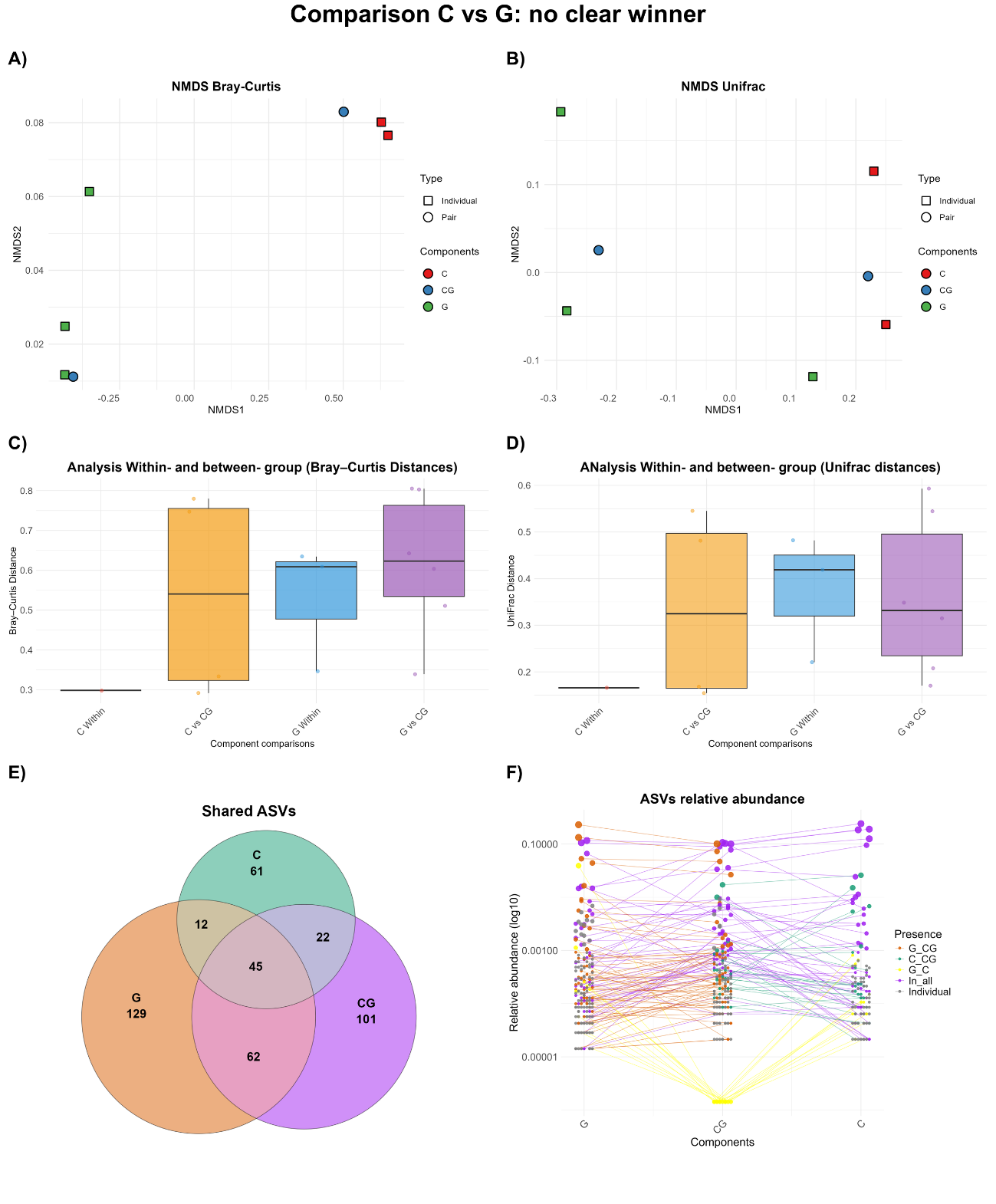

**Supplementary Figure 11**

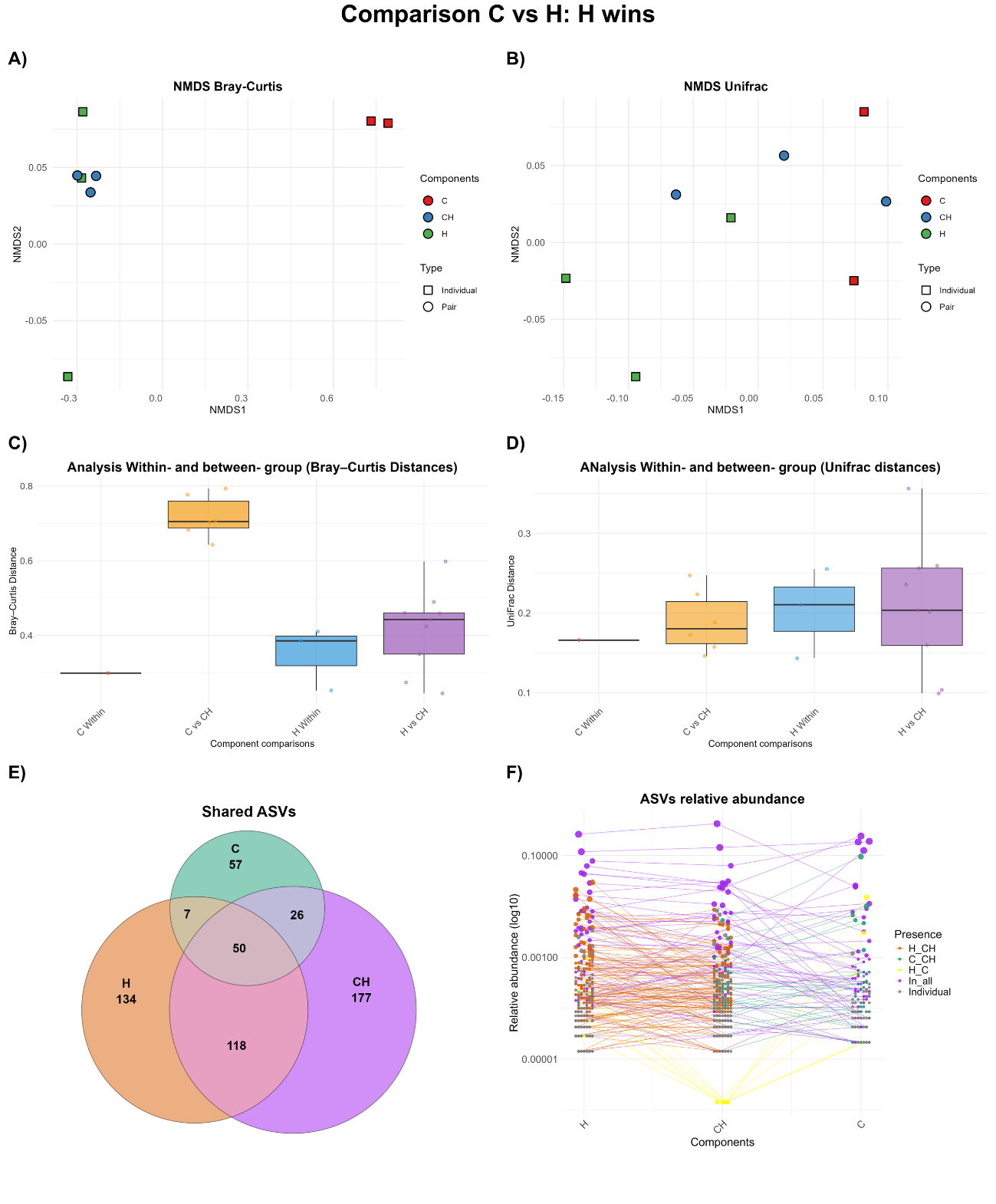

**Supplementary Figure 12**

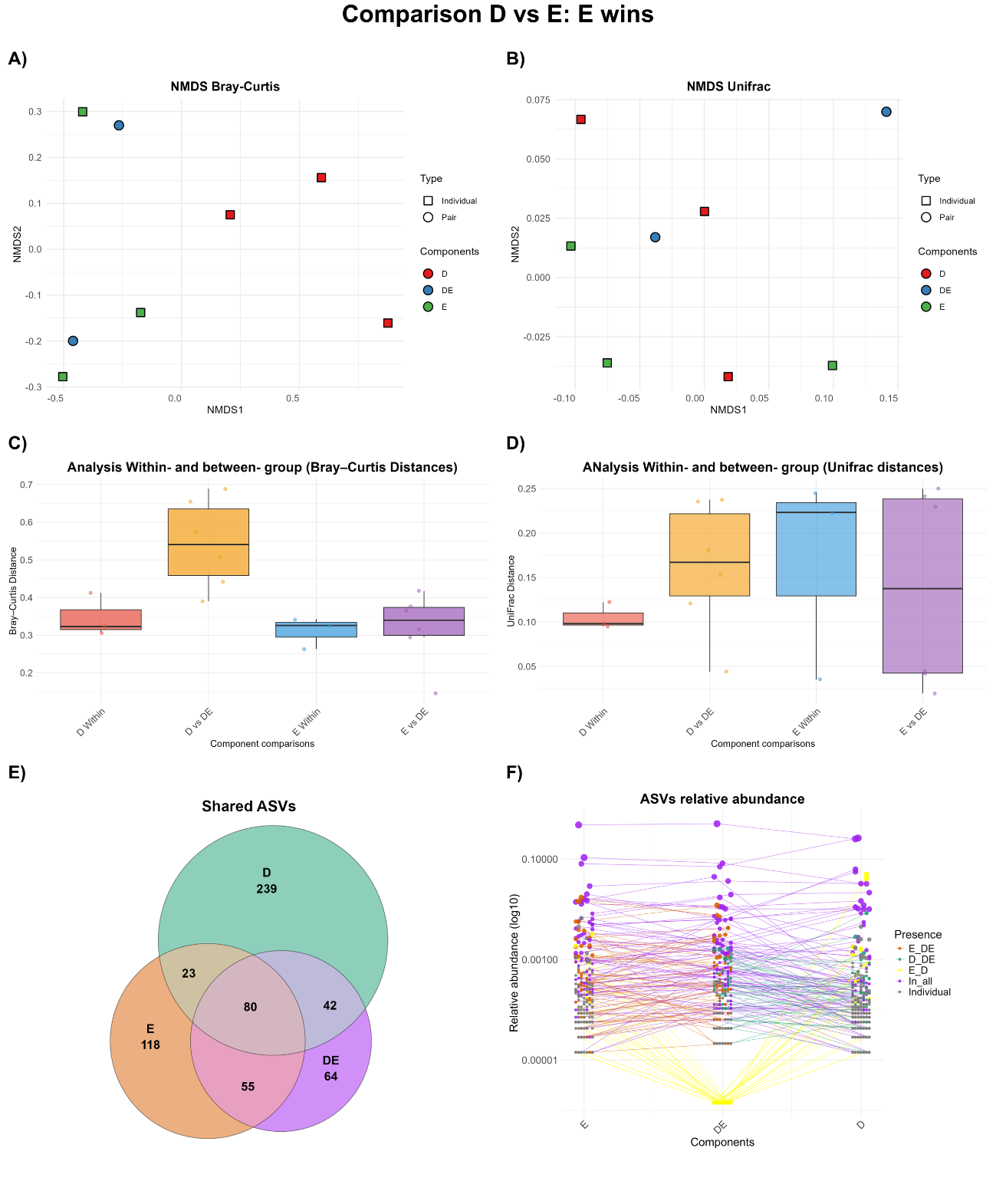

**Supplementary Figure 13**

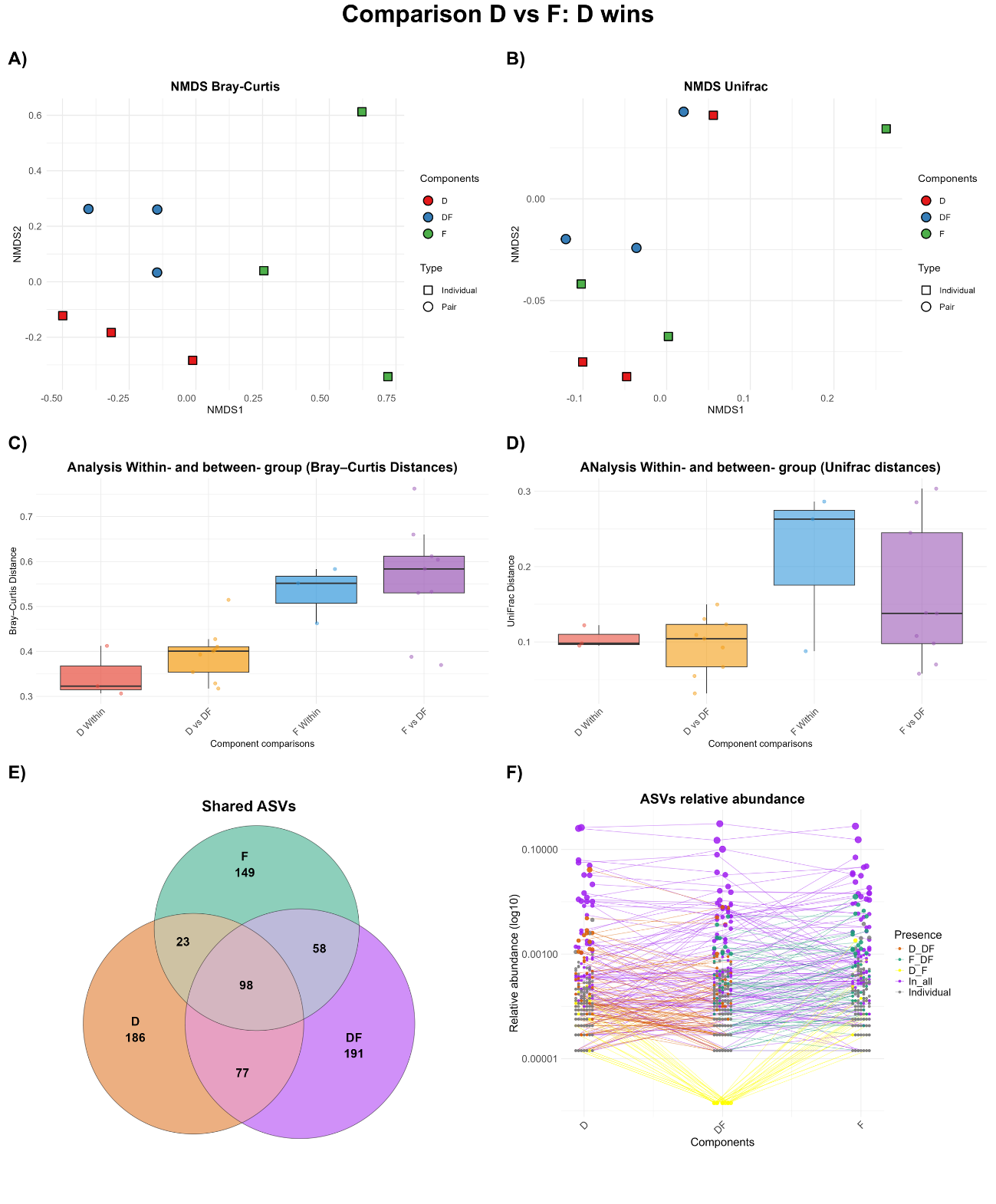

**Supplementary Figure 14**

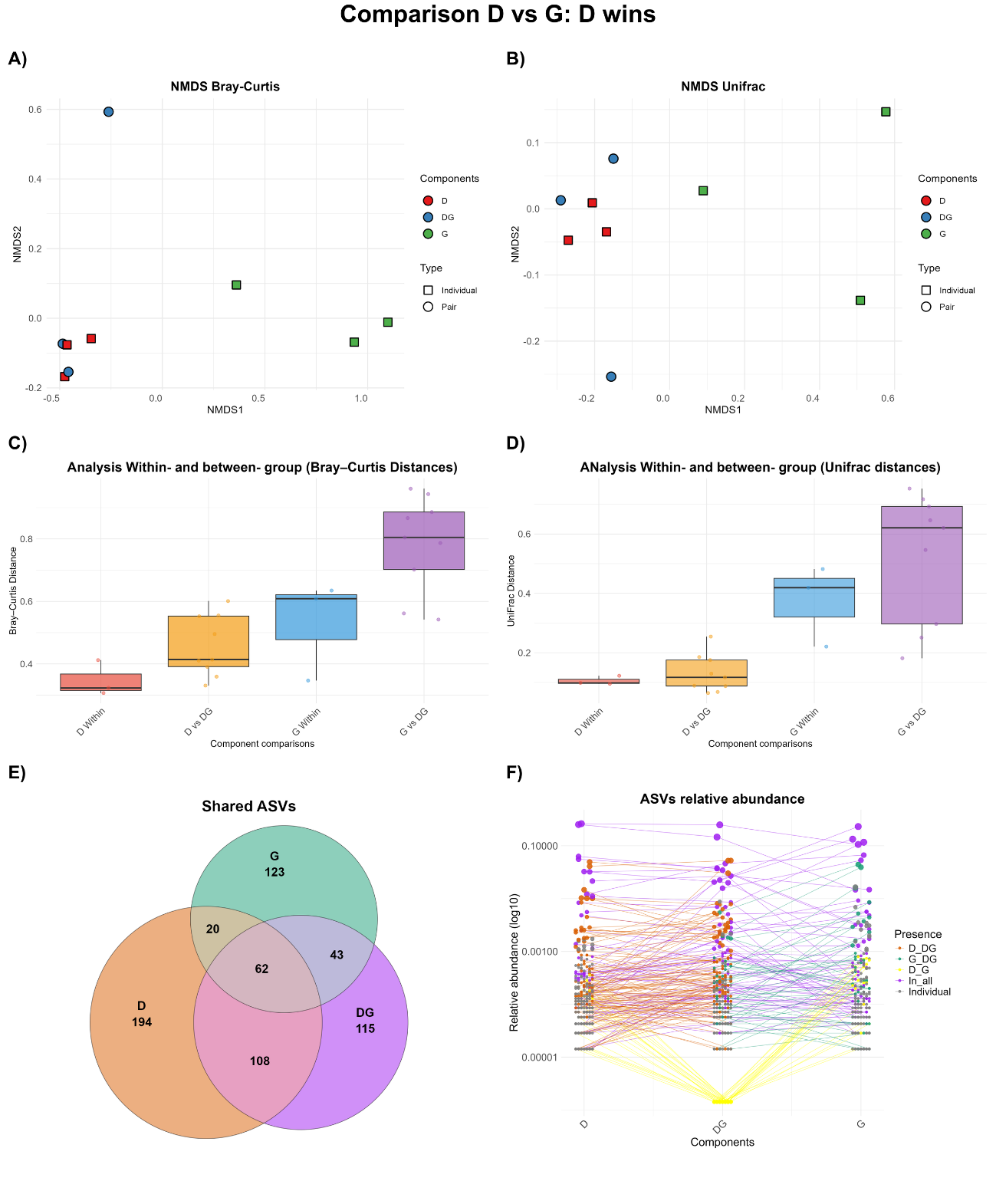

**Supplementary Figure 15**

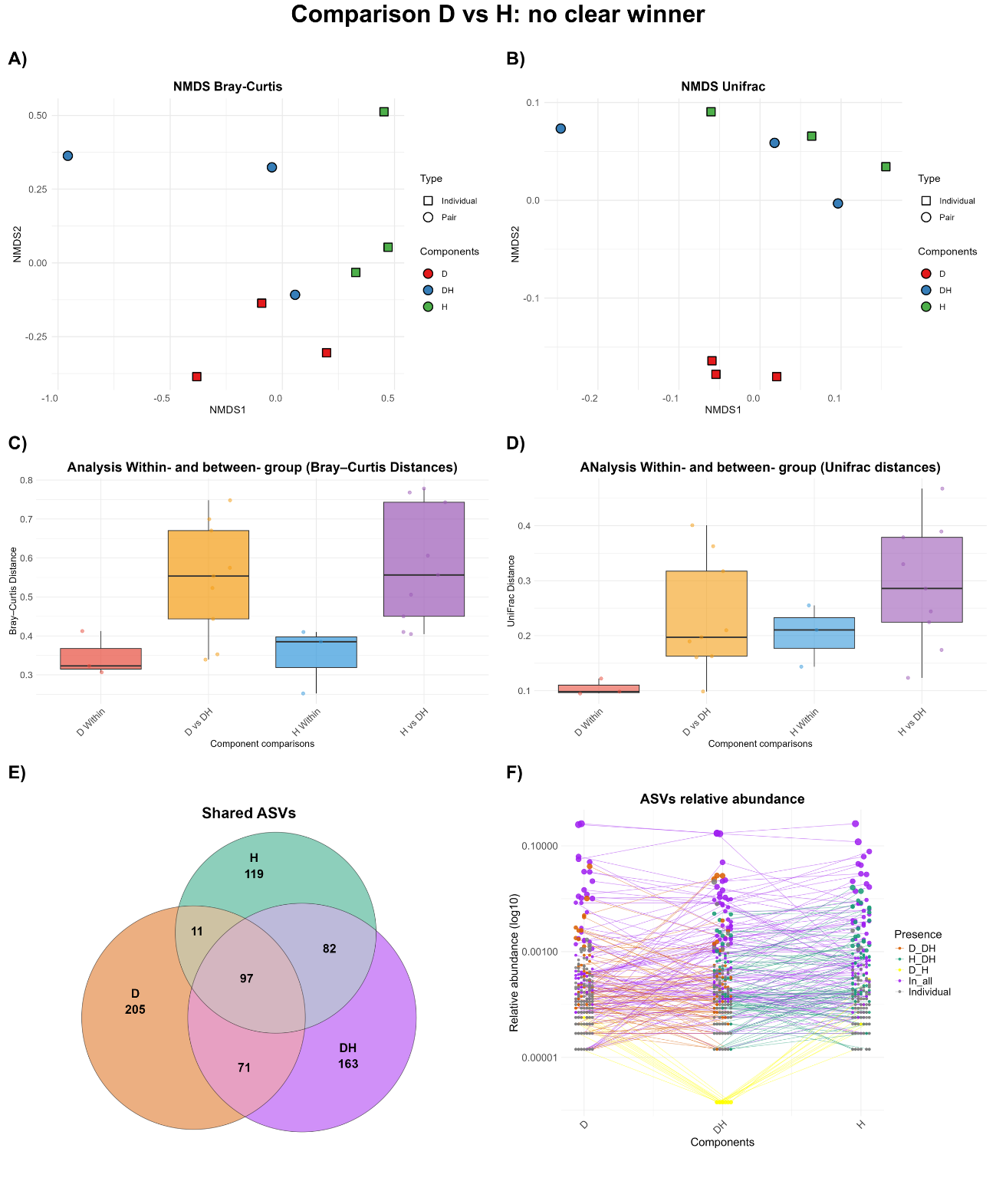

**Supplementary Figure 16**

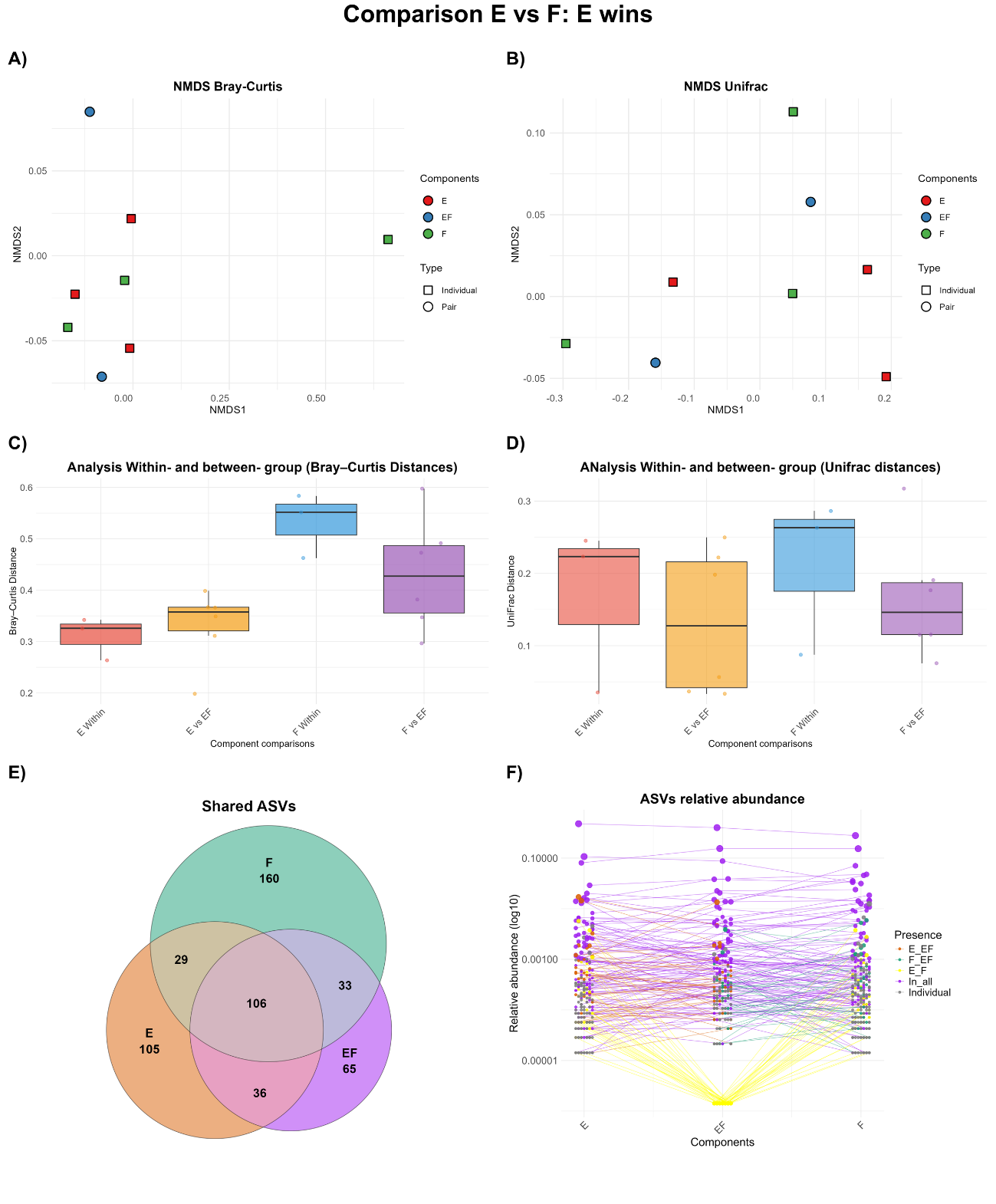

**Supplementary Figure 17**

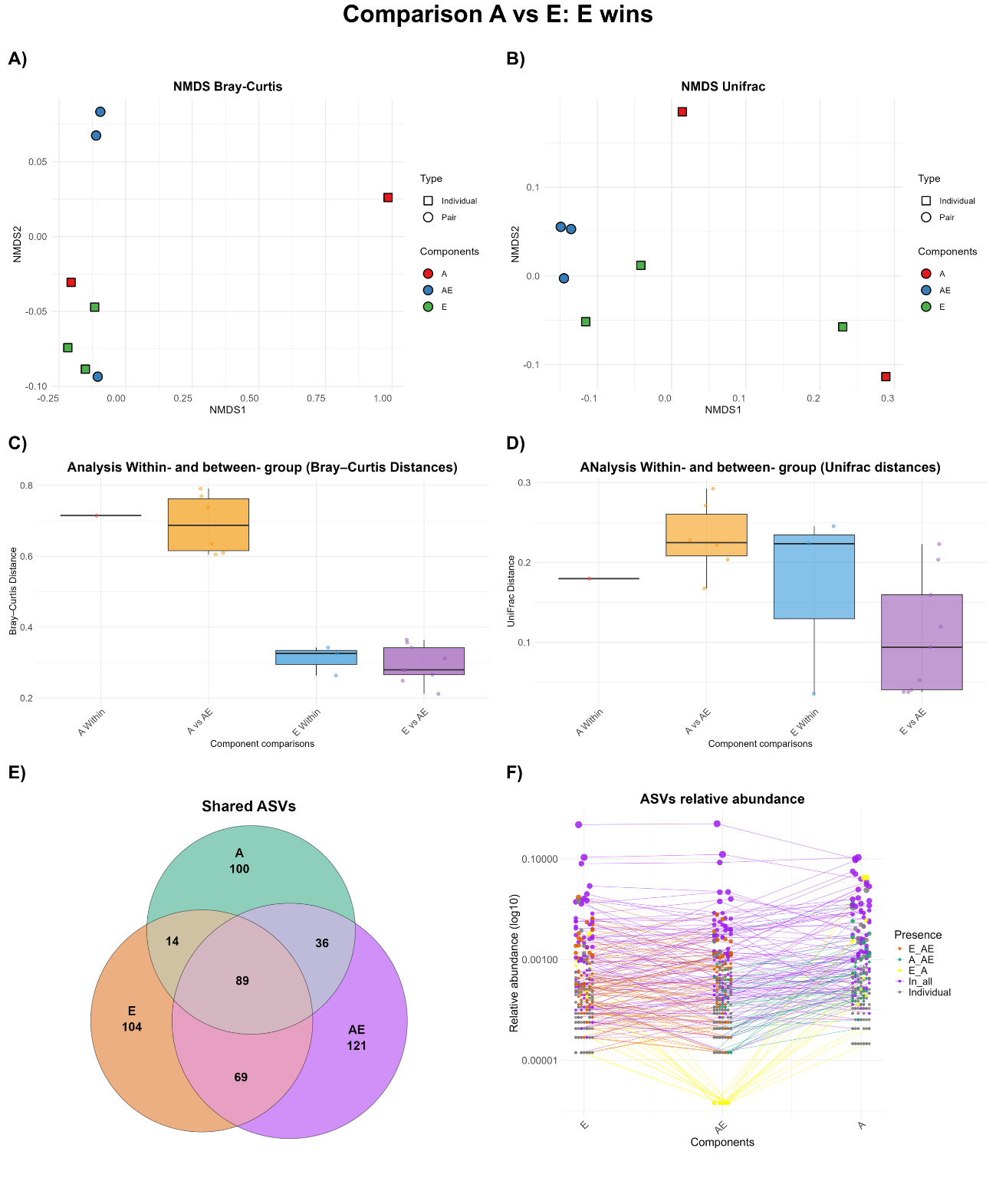

**Supplementary Figure 18**

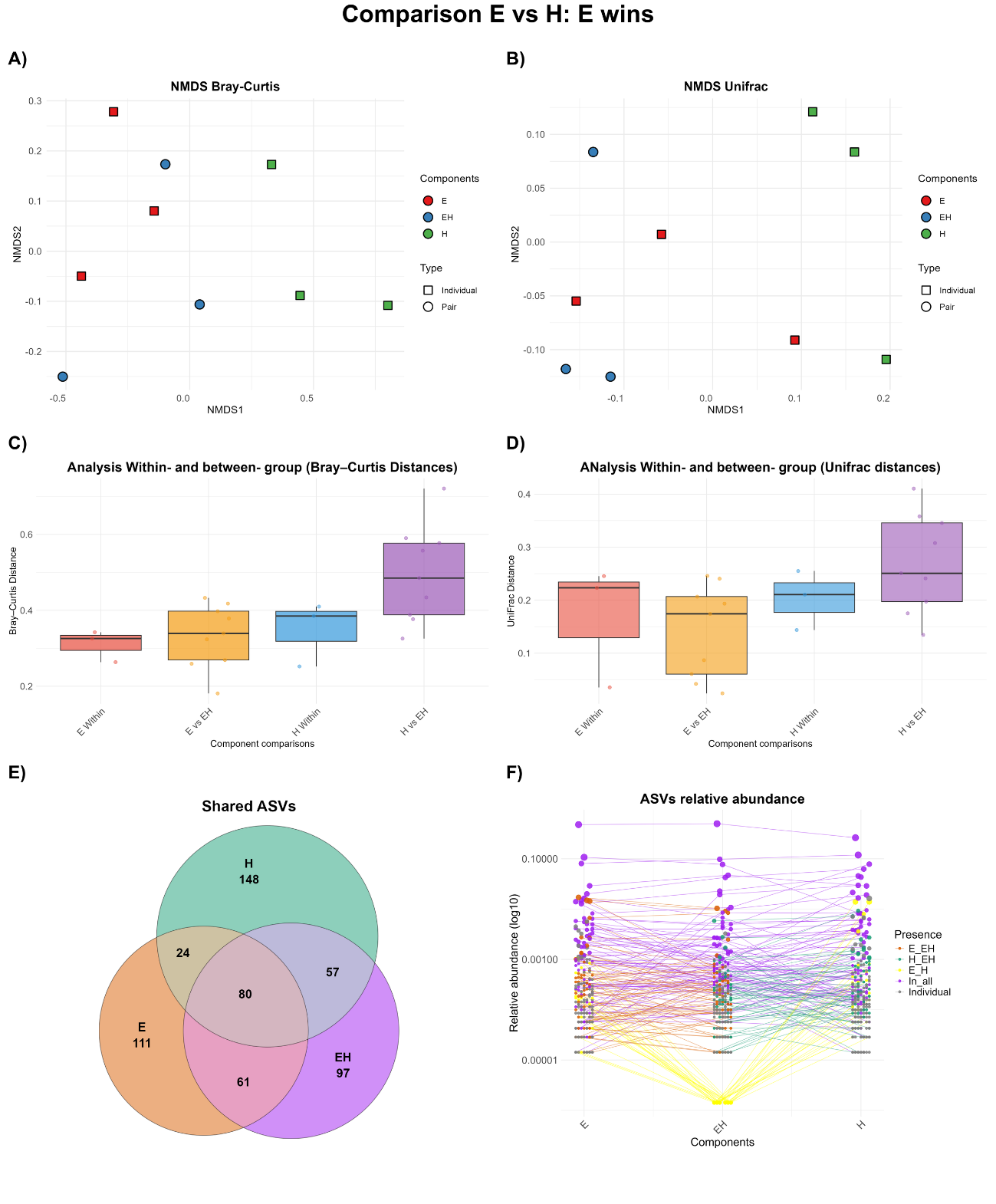

**Supplementary Figure 19**

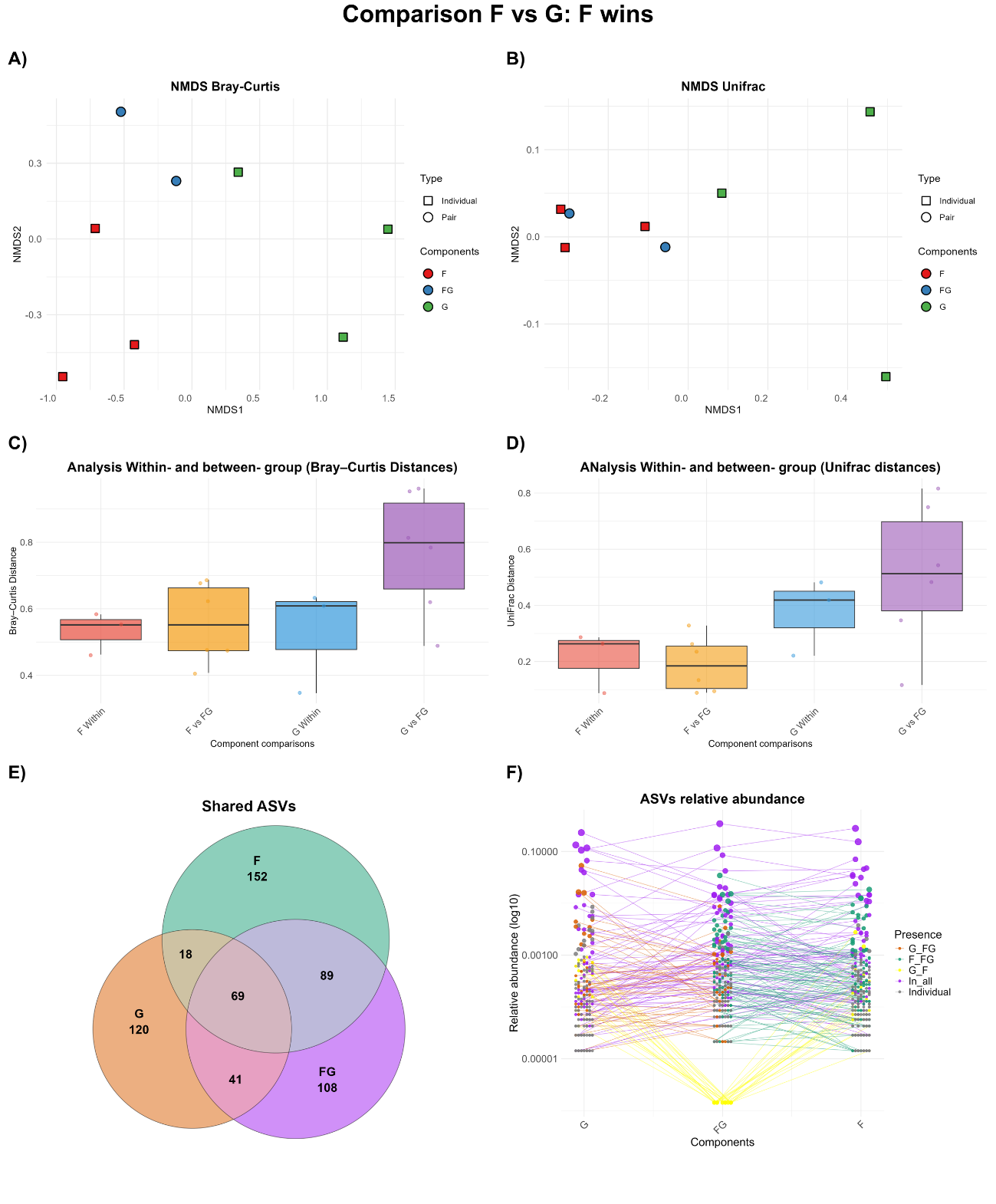

**Supplementary Figure 20**

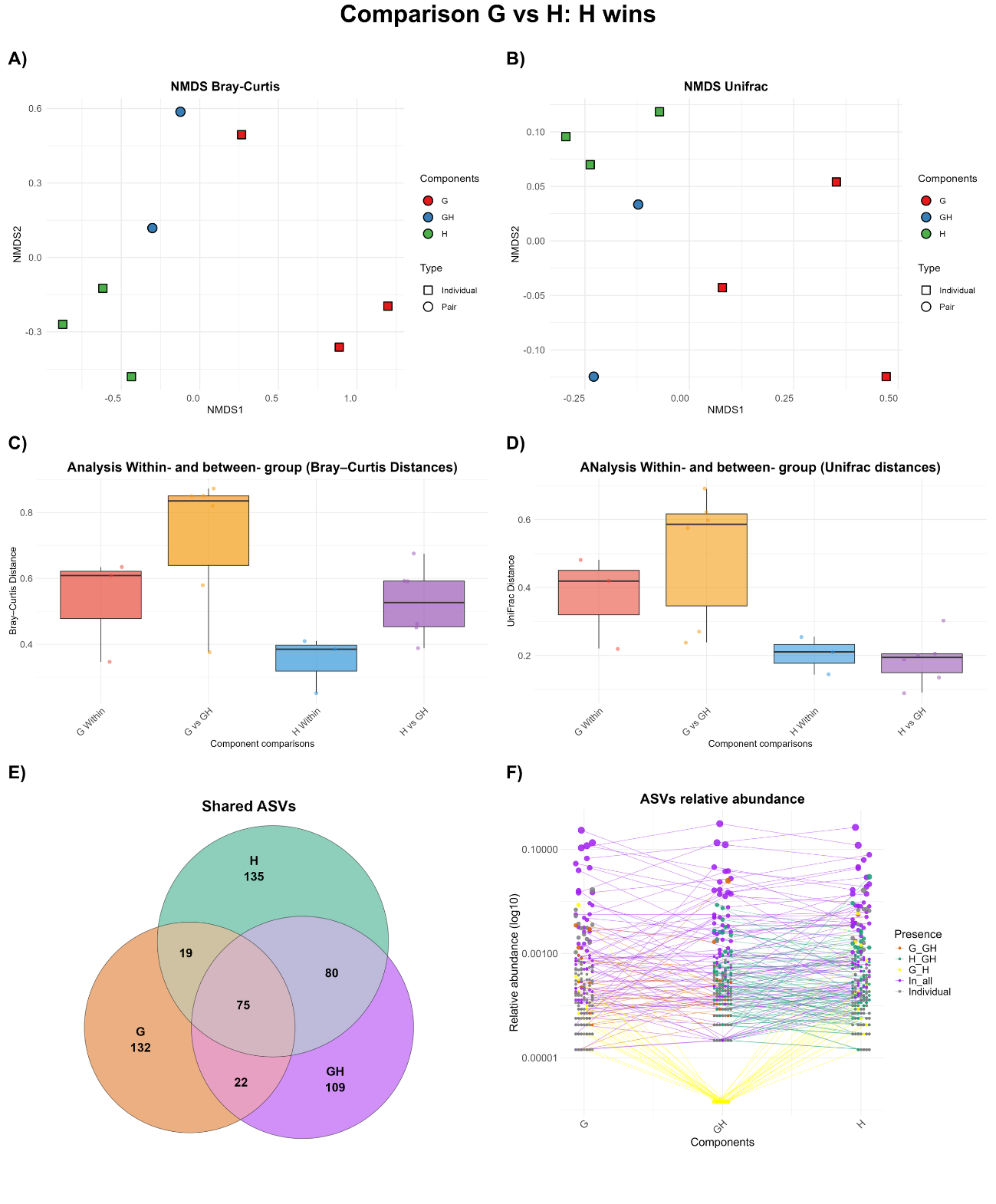

**Supplementary Figure 21**

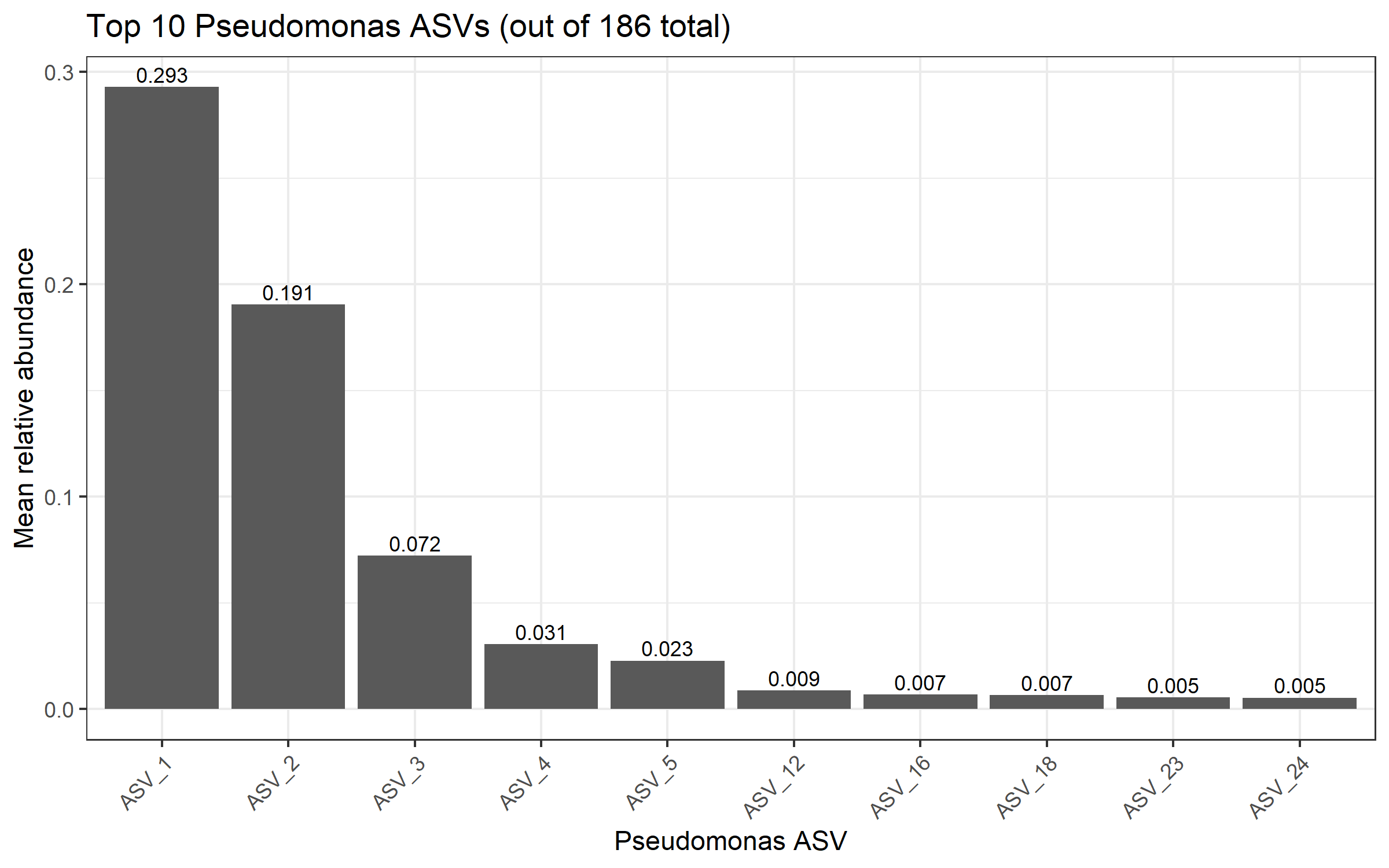

**Supplementary Figure 22**. Mean relative abundance of the 10 most abundant Pseudomonas ASVs across all rhizosphere samples. From a total of 186 ASVs taxonomically assigned to *Pseudomonas*, the 10 most abundant ASVs are shown, ranked by ASV identifier. Bar heights indicate the mean relative abundance of each ASV across all samples, and values above bars indicate the corresponding mean relative abundance. This distribution highlights the strong dominance of a small subset of *Pseudomonas* ASVs within the broader *Pseudomonas* diversity.

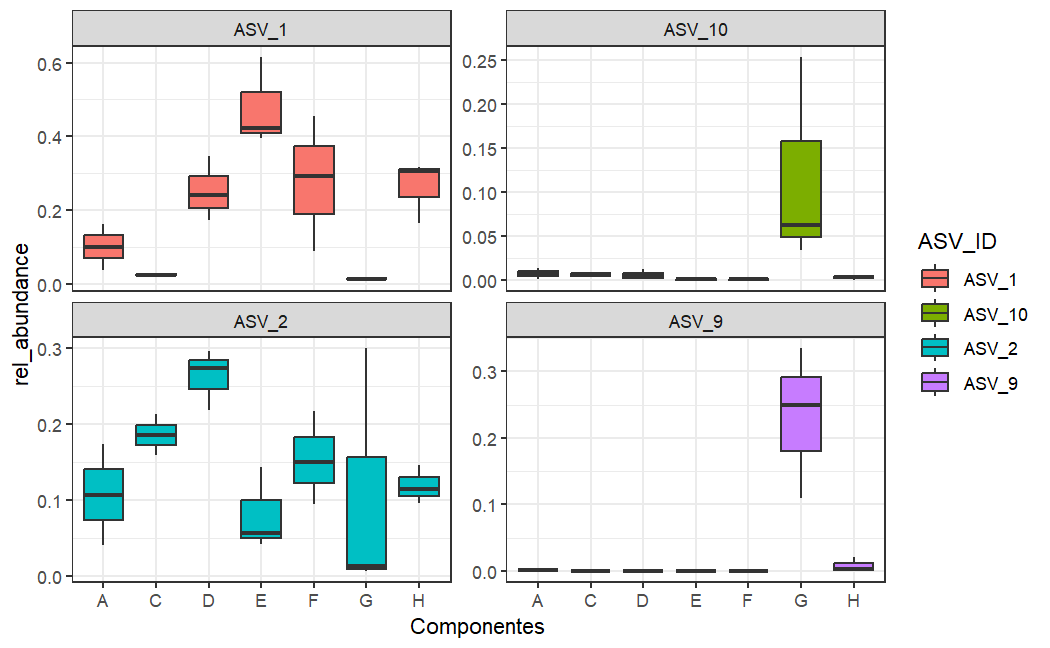

**Supplementary Figure 23**. Boxplots show the distribution of relative abundance for ASV_1, ASV_2, ASV_9, and ASV_10 across samples inoculated with communities A, C, D, E, F, G, and H. Boxes represent the interquartile range (IQR), horizontal lines indicate the median, and whiskers extend to 1.5 × IQR.
